# Interferon-mediated NK cell activation is associated with limited neutralization breadth during SARS-CoV-2 infection

**DOI:** 10.1101/2024.10.22.619639

**Authors:** Izumi de los Rios Kobara, Radeesha Jayewickreme, Madeline J. Lee, Aaron J. Wilk, Stanford COVID-19 Biobank, Andra L. Blomkalns, Kari C. Nadeau, Samuel Yang, Angela J. Rogers, Catherine A. Blish

## Abstract

Best known for their ability to kill infected or malignant cells, natural killer (NK) cells are also underappreciated regulators of the antibody response to viral infection. In mice, NK cells can kill T follicular helper (Tfh) cells, decreasing somatic hypermutation and vaccine responses. Although human NK cell activation correlates with poor vaccine response, the mechanisms of human NK cell regulation of adaptive immunity are poorly understood. We found that in human ancestral SARS-CoV-2 infection, broad neutralizers, who were capable of neutralizing Alpha, Beta, and Delta, had fewer NK cells that expressed inhibitory and immaturity markers whereas NK cells from narrow neutralizers were highly activated and expressed interferon-stimulated genes (ISGs). ISG-mediated activation in NK cells from healthy donors increased cytotoxicity and functional responses to induced Tfh-like cells. This work reveals that NK cell activation and dysregulated inflammation may play a role in poor antibody response to SARS-CoV-2 and opens exciting avenues for designing improved vaccines and adjuvants to target emerging pathogens.

## Introduction

SARS-CoV-2 infection results in mild to severe respiratory illness and has caused over 7 million deaths since the COVID-19 pandemic began in 2020^1,2^. Many individuals infected with ancestral SARS-CoV-2 develop antibodies that neutralize other variants; however, the breadth and potency of antibody repertoires vary across the population^3,4^. We define breadth as the ability of an individual’s antibody repertoire to neutralize multiple viral variants to which the individual has not been exposed. In SARS-CoV-2 infection, the antibody repertoire from infection or vaccination with the ancestral strain (Wu-1) have variable ability to neutralize variants of concern that evolved later in the pandemic such as Alpha, Beta, Delta, and Omicron. Factors that influence antibody breadth, particularly against variants which the individual has not been exposed to are poorly understood. Here, we sought to identify features of the host peripheral immune response to ancestral SARS-CoV-2 infection that correlate with antibody breadth against future variants to inform our understanding of how to develop improved vaccines against coronaviruses and other pathogens.

Natural killer (NK) cells are understudied regulators of antibody responses^5–16^. Classically known as innate lymphocytes that kill infected or malignant cells, NK cells also play immunoregulatory roles during viral infection due to their interactions with other immune cells^12^. In mice, NK cells can kill T follicular helper (Tfh) cells in the germinal center which limits the level of somatic hypermutation (SHM) and decreases the quality of the antibody response in the setting of vaccination^10^. *In vitro*, NK cells can kill activated CD8 T cells^17–19^, CD4 T cells^10,11,13,20^, and dendritic cells^21,22^ and directly interact with B cells^23,24^. Human clinical trials for yellow fever, malaria, and hepatitis B vaccines have identified NK cells as correlates of non-response. In each trial, higher antibody titers or protection from live pathogen challenge was associated with lower NK cell activation or reduced expression of NK cell functional gene modules^8,9,16^. In the context of infection, dysfunctional NK cells are associated with broadly neutralizing antibodies in individuals with untreated, chronic HIV infection^25^. NK cells can also play a positive role in the antibody response; NK cell secretion of IFNγ is critical to the efficacy of the AS01 adjuvant in mice^7^. These multifaceted roles of NK cells in controlling adaptive immunity remain understudied in both vaccination and infection and represent an important gap in our understanding of human antibody responses.

In order to determine if NK cell phenotype correlates with antibody breadth in SARS-CoV-2 infection, we utilized publicly available data collected by our lab in March - June 2020. This dataset contains profiling of COVID-19 participants across the severity spectrum by both single cell RNA sequencing (scRNA-seq) and mass cytometry by time of flight (CyTOF)^26,27^. For this study, matched serum obtained from the Stanford COVID-19 biobank was used to evaluate the antibody repertoire, enabling us to evaluate immune correlates of breadth in individuals from across the severity spectrum. We examined systemic immune differences between broad and narrow neutralizers and found major differences in the NK cell compartment. NK cells from narrow neutralizers were highly activated and expressed markers of cytotoxicity at both the mRNA and protein levels, including many interferon-stimulated genes (ISGs). Finally, we validated *in vitro* that ISG-driven activation in NK cells resulted in greater functional responses and killing of induced T follicular helper-like (iTfh-like) cells. Overall, this study demonstrates that NK cell phenotype is highly correlated with neutralization breadth in COVID-19 and suggests that ISG-driven inflammation may contribute to narrow neutralization.

## Results

### Cohort Description

Our lab previously profiled 30 COVID-19 participants from the entire severity spectrum and 7 healthy controls by scRNA-seq as well as 26 participants (a subset of the 30 in scRNA-seq) and 12 healthy controls by CyTOF^26,27^ (Figure 1A). This cohort was well-controlled for both viral variant and prior exposure to SARS-CoV-2 because these samples were collected during the first 4 months of the COVID-19 pandemic. Thus, all individuals experienced primary infection with ancestral SARS-CoV-2 because variants of concern had not yet evolved^4^. PBMCs and red blood cell-lysed whole blood were profiled by scRNA-seq, and PBMCs as well as untouched, magnetically-purified natural killer (NK) cells were profiled by CyTOF. This cohort contains COVID-19 participants from across the full range of the WHO disease severity spectrum, including participants with mild disease who remained un-hospitalized (WHO 1-3), hospitalized participants with moderate disease (WHO 4-5), hospitalized participants who required intubation or ECMO (WHO 6-7), and participants who were later deceased (WHO 8). Samples were collected 0 - 66 days post positive COVID-19 test (median 4 days) and the majority of samples were drawn during the acute phase of disease (Extended Data Figure 1D).

**Figure 1:**
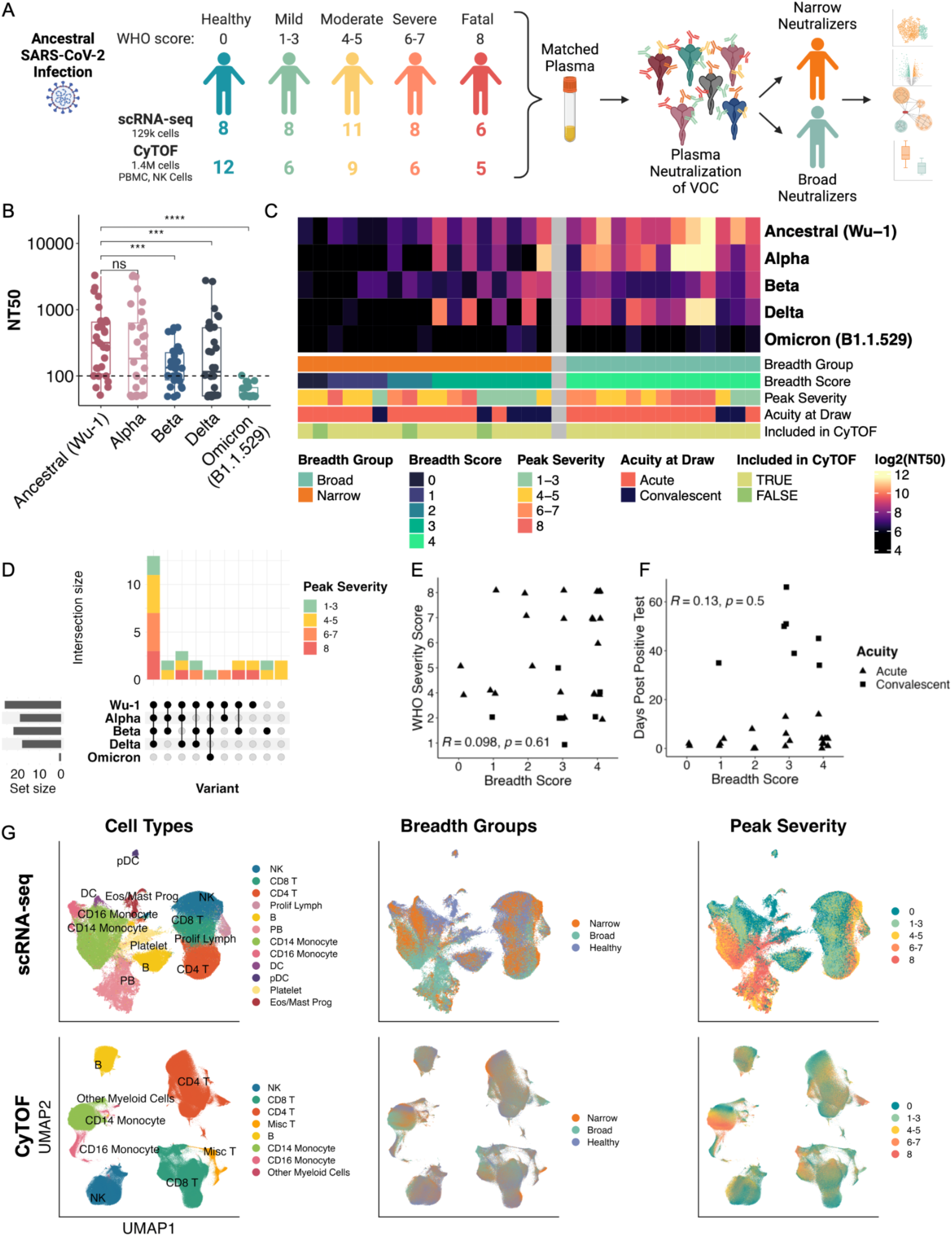
Heterogeneity in Neutralization Breadth against Variants of Concern after Ancestral SARS-CoV-2 Infection. **A.** Study outline of number of samples profiled, summarized by modality and peak WHO severity score and analysis plan. **B.** 50% Neutralization titers (NT50) and median titer against 5 SARS-CoV-2 variant pseudoviruses. *, P < 0.05; **, P < 0.01; ***, P < 0.001; ****, P < 0.0001; ns, P > 0.05 by two-sided, paired Wilcoxon rank-sum test with Bonferroni’s correction for multiple hypothesis testing Each point represents one donor and the average of 3 technical replicates. **C.** Heatmap of NT50 against 5 variants of SARS-CoV-2 pseudovirus for each patient. Each cell represents the average of 3 technical replicates. D. UpSet plot of all unique combinations of SARS-COV-2 variants neutralized by each patient colored by peak WHO Severity Scores. **E.** Scatter plot of breadth score and WHO severity score. **F.** Scatter plot of breadth score and days post positive test. P-values by Spearman rank correlation **G.** UMAP projections of all PBMCs of complete scRNA-seq (top) and CyTOF (bottom) datasets colored by cell types, breadth groups, and peak WHO severity score. PB = plasmablast, Eos = eosinophil; Prog = progenitor; Prolif Lymph = proliferating lymphocyte.

### Variability in antibody breadth during ancestral-SARS-CoV-2 infection is independent of severity

To determine immunological correlates of antibody breadth in SARS-CoV-2 infection, we evaluated antibody neutralization breadth against variants of concern using matched plasma from the Stanford COVID-19 Biobank for each corresponding COVID-19 patient with publicly available scRNA-seq and CyTOF data. Matched plasma samples were from the same or closest blood draw (one donor’s closest plasma sample was +3 days and remainder were from the same draw). All individuals were infected with ancestral SARS-CoV-2, equalizing our measurement of breadth against variants of concern. To stratify participants by breadth against variants of concern, we measured serum neutralization against either ancestral SARS-CoV-2 (Wu-1) pseudovirus or one of 4 variants which emerged after patient samples were collected (Alpha, Beta, Delta, and original Omicron B.1.1.529) in a HeLa cell line stably expressing ACE2 and TMPRSS2^28^ (Figure 1 A-D). We found that there was heterogeneity in both the neutralization potency against each variant and in the number of variants neutralized (Figure 1B-D). Median NT50 (the reciprocal of the highest dilution to obtain <50% infection) against Wu-1 was 317 with about 1 log of variation in NT50 across individuals. Median NT50 was significantly reduced from Wu-1 in Beta (133), Delta (117), and Omicron B.1.1.529 (50), and NT50 against these variants had a broad range across the cohort (Figure 1B). As our cohort showed little neutralizing activity against original Omicron, we only included the first 4 major variants. Using the number of variants neutralized above the limit of detection as a breadth score, participants were ranked by breadth (Figure 1B-D). Participants neutralized between 0 and 4 variants; 13 participants neutralized Wu-1, Alpha, Beta, and Delta and were classified as broad, while those that neutralized different combinations of 0-3 variants were classified as narrow (Figure 1C and D).

As it is possible that severity could drive breadth via immune activation, we validated that breadth score did not correlate with WHO severity score (R = 0.098, P = 0.61) (Figure 1E). Severity score was also well-distributed over different variant neutralization combinations (Figure 1D). Therefore, we can be confident that immunological signatures of breadth cannot be primarily attributed to severity, and, by including participants across the entire severity spectrum, we can analyze immune correlates of antibody breadth regardless of immunological signatures of disease severity. Additionally, sample collection day post-positive test did not correlate with breadth scores (R = 0.13, P = 0.5), so breadth cannot be primarily attributed to acute or convalescent sample identity (Figure 1F). Breadth score and breadth groups were assigned to each cell. In UMAP projections of scRNA-seq and CyTOF data, there is representation of major immune cell types in both broad and narrow neutralizers, allowing us to thoroughly investigate systemic contributions to a productive immune response with regard to antibody breadth (Figure 1G).

### Natural killer cells in narrow neutralizers of SARS-CoV-2 highly express interferon-stimulated genes and markers of activation

To identify the most robust immunological signal correlated with breadth, we used pseudobulk whole PBMC samples from each patient to identify genes that significantly correlated with breadth score using DESeq2^29^. In whole PBMCs, we identified 25 genes whose expression significantly correlated with low breadth score, 24 of which were interferon-stimulated genes (ISGs) verified by the interferome database including classical ISGs like MX family, OAS family, and IFI family genes^30^ (Figure 2A). We next examined which cell types in narrow and broad breadth groups expressed these genes and found that NK cells from narrow neutralizers were amongst the highest expressors of the genes correlated with low neutralization score (Figure 2B). We also found NK cells to be one of only two cell types with differences in cell type proportion between broad and narrow neutralizers. NK cells as a proportion of PBMCs are inversely correlated with breadth score in both scRNA-seq (R = −0.47, P = 0.0092) and CyTOF modalities (R = −0.48, P = 0.012); this trend was also observed in breadth groups where broad neutralizers have significantly less NK cells when compared to narrow neutralizers and healthy controls (median % of NK cells in scRNA-seq: broad = 5.97, narrow = 12.6, healthy = 20.2; CyTOF: broad = 8.68, narrow = 14.0, healthy = 19.3) (Figure 2C-F). pDCs (<1% of PBMCs in most participants) were significantly more abundant in narrow neutralizers in scRNA-seq data (Extended Data Figure 2A-B). This perturbation of NK cells between breadth groups and breadth scores motivated us to further analyze the role of the NK cell compartment in modulating antibody breadth.

**Figure 2:**
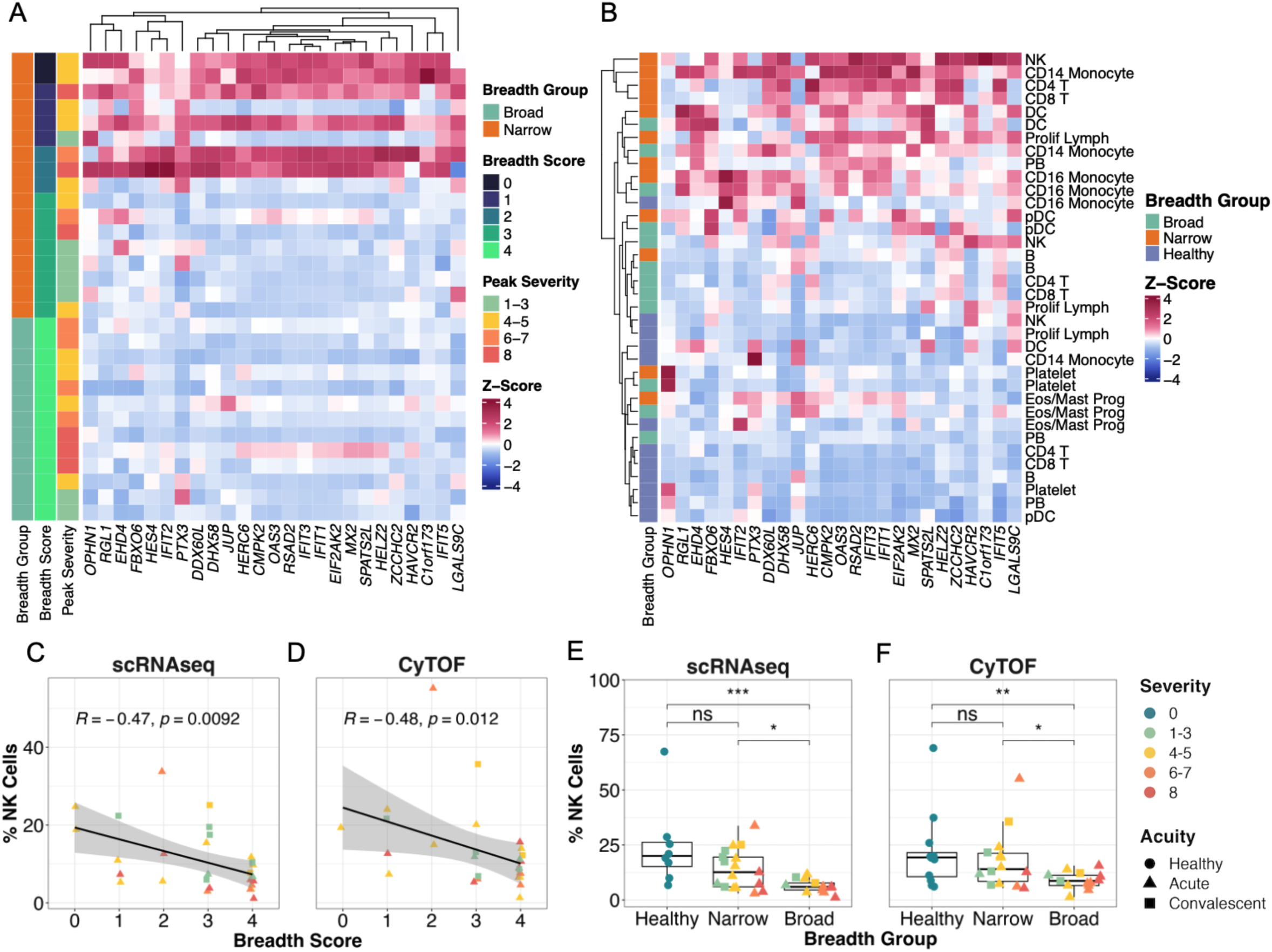
Interferon-Stimulated Genes are Associated with Narrow Neutralizers and are Highly Expressed in NK cells. **A.** Heatmap of Z-score and count-normalized RNA expression of genes significantly correlated with low neutralization score in all PBMCs from DESeq2 pseudobulk correlation analysis. Genes (columns) ordered by hierarchical clustering. **B.** Heatmap of Z-score normalized expression of interferon-stimulated genes from DESeq2 pseudobulk analysis for each cell type and breadth group ordered by hierarchical clustering for rows. **C-D**. Correlation between proportion of NK cells and breadth score in scRNA-seq (C) and CyTOF(D) datasets. P-value by Spearman rank correlation. **E-F.** Boxplots of proportion of NK cells in each breadth group from scRNA-seq (E) and CyTOF (F) datasets. *, P < 0.05; **, P < 0.01; ***, P < 0.001; ns, P = 0.05 by two-sided Wilcoxon rank-sum test with Bonferroni’s correction for multiple hypothesis testing.

In UMAP space, we observed separation of NK cells by both severity and breadth groups (Figure 3A and B). We identified differentially expressed genes (DEGs) in NK cells between broad and narrow breadth groups (log_2_FC > 0.25 and adjusted P-value < 0.05). NK cells from narrow neutralizers express multiple ISGs such as *IFI44L*, *RSAD2*, *XAF1*, *IFIT3*, *MX1*, *IFIT1* etc. and perforin (*PRF1*), a marker of NK cell cytotoxicity (Figure 3C, Table 3). NK cells from narrow neutralizers also upregulated *CX3CR1,* which is a marker of migration to lymphoid tissues where NK cells could influence antibody responses as well as associated with cytotoxic function in NK cells^31^.

**Figure 3:**
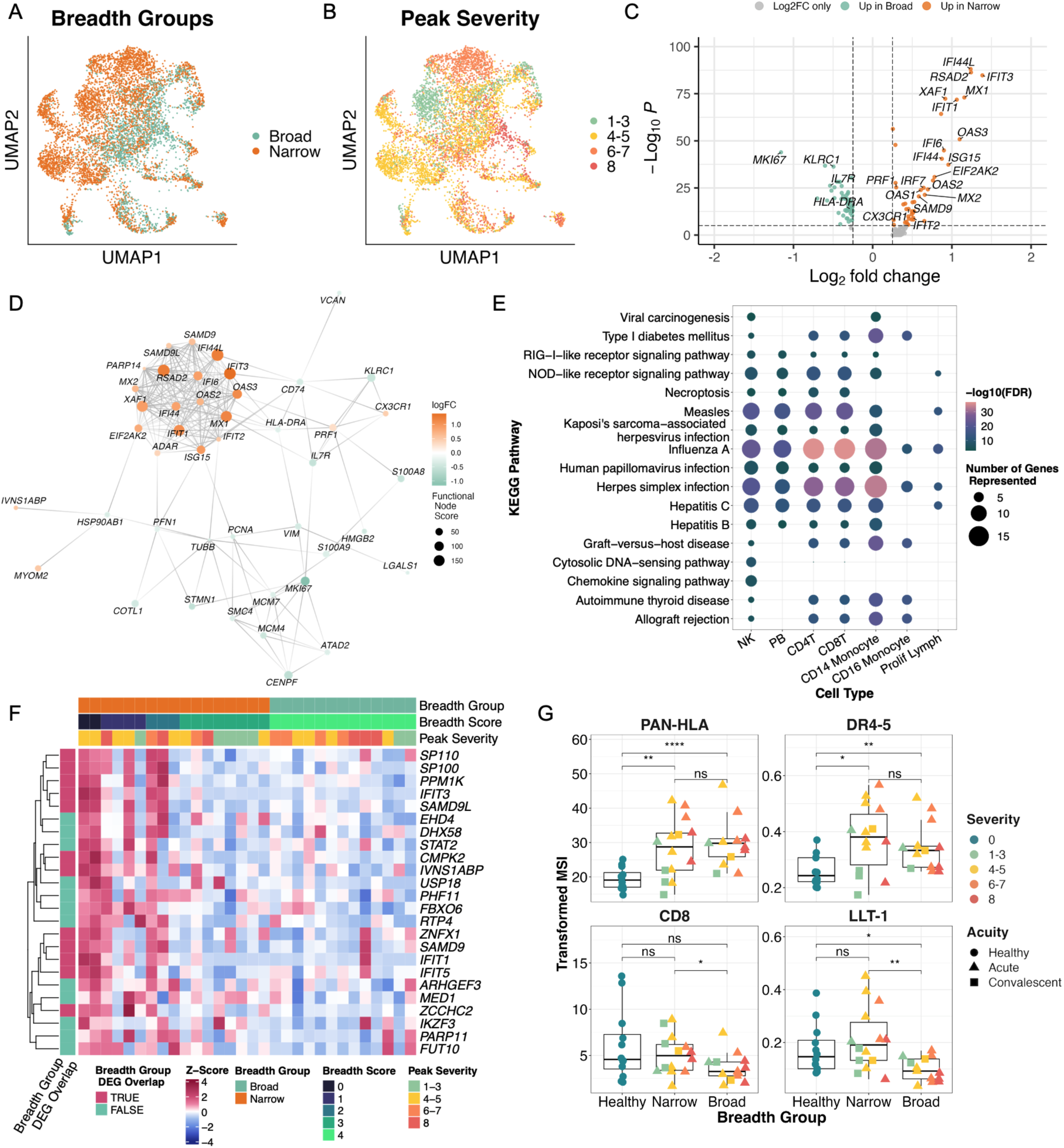
Interferon-Stimulated Genes in Abundant NK cells Correlate with Low Neutralization Breadth. **A-B**. UMAP Projections of NK Cells from scRNAseq dataset colored by breadth group (A) or peak WHO severity score (B). **C.** Volcano plot of differentially expressed genes between broad and narrow neutralizers from NK cells in scRNAseq dataset. P-value by Wilcoxon rank-sum test with Bonferroni adjustment for multiple hypotheses. **D**. Protein-protein interaction graph depicting highest-scoring, minimal significant interacting graph of differentially expressed genes in NK cells from COVID-19. **E.** Dot plot depicting significantly enriched KEGG pathways colored by false discovery rate (FDR) from pathway analysis (see methods), sized by number of genes in KEGG pathway present in each cell type, and ordered by hierarchical clustering. Pathways were chosen by significant enrichment in NK Cells from narrow neutralizers. **F.** Heatmap of Z-score and count normalized transcript level expression of genes significantly correlated with low neutralization score from DESeq2 pseudobulk correlation analysis in NK cells from COVID-19 participants. **G.** Boxplots quantifying arcsinh-transformed average expression of markers from CyTOF dataset. *, P < 0.05; **, P < 0.01; ***, P < 0.001; ****, P < 0.0001; ns, P = 0.05 by two-sided Wilcoxon rank-sum test with Bonferroni’s correction for multiple hypothesis testing.

In order to visualize the interaction landscape and underlying regulators of DEGs in NK cells, we leveraged the Bionet package to find the highest-scoring subgraph of DEGs among a known protein-protein interaction network derived from the human STRING database^32^. Using DEGs between broad and narrow neutralizers in NK cells, we constructed a minimal significant network (FDR < 1e-6) of protein-protein interactions from the background interaction graph of the 2000 most variable genes in NK cells from COVID-19 participants. Nodes are then scored using a β-uniform mixture model fitted to the adjusted P-value distribution^33,34^. The minimal significant network of DEGs from NK cells revealed a hub of highly differentially-expressed and interacting ISGs (*MX1, IFIT1, ISG15* etc.) in narrow neutralizers with high predicted functional node scores relative to other nodes (Figure 3D). This confirms a high level of ISG-mediated activation in NK cells from narrow neutralizers. *CX3CR1* and Perforin (*PRF1*) were also included in the minimally significant graph, highlighting both the migratory and cytotoxic potential of NK cells from narrow neutralizers.

Using DEGs upregulated in narrow neutralizers across all cell types, we performed pathway analysis to identify KEGG pathways enriched in narrow neutralizers. NK cells from narrow neutralizers were enriched for multiple pathways related to viral infections, innate immune sensing, and pathological inflammation (e.g. Influenza, Cytosolic DNA sensing, and NOD/RIG-I signaling pathways). Many of these pathways were also significantly enriched in multiple other immune cell types in narrow neutralizers (Figure 3E). This indicates narrow neutralizers have heightened inflammation across the immune system.

We also repeated our pseudobulk correlation analysis using DESeq2 to find genes significantly correlated with breadth score in NK cells (Figure 3F). We found that all genes that were significantly correlated with low breadth score were ISGs (verified by interferome database)^30^; the majority overlapped with scRNA-seq DEGs from breadth groups (Figure 3C and F). On the protein level, all COVID-19 participants exhibited upregulation of HLA and death receptors (DR4-5); however, only narrow neutralizers expressed significantly greater levels of CD8 and LLT-1 (markers of cytotoxicity and activation, respectively) when compared to broad neutralizers^35,36^ (Figure 3G). This reveals specific activation markers present only in narrow neutralizer NK cells in the setting of other signals of disease-driven activation.

### NK cells are immature and proliferating in broad neutralizers of SARS-CoV-2

Distinct from narrow neutralizers, in broad neutralizers, NK cells were significantly less abundant, but they express higher transcript levels of *MKI67* (Ki-67), *KLRC1* (NKG2A), and *IL7R* (CD127) indicating a proliferative, inhibitory, and immature phenotype^37,38^ (Figure 3C, Figure 4 A-B). Differential expression analysis indicates upregulation of some genes involved in immune activation like *HLA-DRA*, but no evidence of canonical ISG activation (Figure 3C). This was substantiated on the protein level where NK cells from broad neutralizers expressed significantly higher levels of NKG2A as well as its signaling partner CD94 which is necessary for NKG2A’s inhibitory function (Figure 4A-B). Significantly higher expression of *IL7R* in scRNA-seq space along with elevated levels of CD56 in CyTOF space further confirms the immature and inhibitory NK phenotype in broad neutralizers^39^ (Fig 4A). HLA-DR was upregulated on the protein, but not transcript (aggregated expression of *HLA-DRA*, and *HLA-DRB1-5*) level. Similarly, *KLRD1* (CD94) and *NCAM1* (CD56) were not differentially expressed in scRNA-seq data, but can have poor correlation between transcript and protein expression^40^ (Figure 4 A-B). Additionally, HLA-E, the ligand for NKG2A/CD94 heterodimer, is not differentially expressed on any cell type between broad and narrow neutralizers at the protein level^41^ (Extended Data Figure 3B).

**Figure 4:**
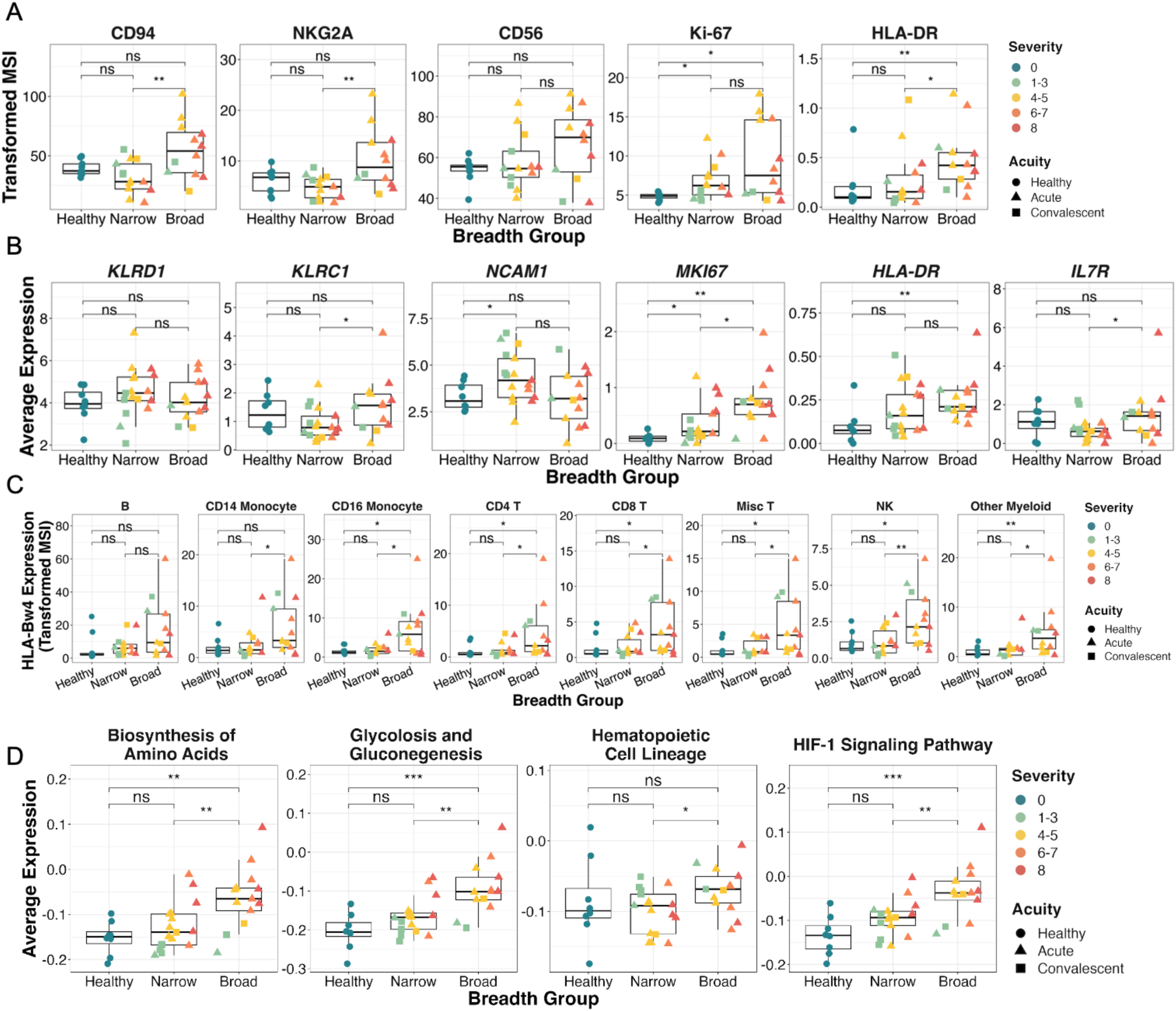
NK Cells From Broad Neutralizers are immature, proliferative and enriched for normal processes. **A**. Boxplots quantifying arcsinh-transformed average expression of markers from CyTOF dataset. **B**. Boxplots quantifying average expression of markers from A and *IL7R* in scRNAseq dataset. HLA-DR is aggregated expression of HLA-DR genes *HLA-DRA* and *HLA-DRB1-5* **C**. Boxplots quantifying arcsinh-transformed average expression of HLA-bw4 in each cell type from CyTOF dataset. **D.** Boxplots quantifying average expression of KEGG module scores in scRNAseq expression data. *, P < 0.05; **, P < 0.01; ***, P < 0.001; ****, P < 0.0001; ns, P = 0.05 by two-sided Wilcoxon rank-sum test with Bonferroni’s correction for multiple hypothesis testing.

All cell types in broad neutralizers express higher levels of HLA-Bw4 (significant in all cell types except B cells) whereas expression of HLA-Bw6 is not different between broad and narrow neutralizers (Figure 4C and Extended Data Figure 3A). HLA-Bw4 and HLA-Bw6 represent mutually exclusive epitopes on class I HLA-B and HLA-Bw4 but not HLA-Bw6 is directly recognized by NK cells and is correlated with protection from COVID-19 and control of HIV^42^. This may support the partial contributions of genetics to productive antibody response to SARS-CoV-2. A KIR3DL1+HLA-Bw4+ genotype is also reported to be associated with protection from severe COVID-19^43^; however, neither KIR3DL1 nor any other KIR measured or detected in CyTOF or scRNA-seq data was differentially expressed between any group (Extended Data Figure 3C and D).

Using genes significantly upregulated at the RNA level in NK cells from broad neutralizers, we performed pathway analysis using the KEGG pathway database^44^. We found upregulation of multiple pathways indicating normal cell functions and immaturity in NK cells from broad neutralizers, including Biosynthesis of amino acids, Glycolysis and Gluconeogenesis, Hematopoietic Cell Lineage, and HIF-1 Signaling (Figure 4D, Table 3).

### NK cell transcriptomic clusters enriched in narrow neutralizers express cytotoxic proteins and differentiation markers

To further investigate the phenotype of NK cells enriched in broad and narrow neutralizers, we applied unsupervised clustering to scRNA-seq data of all NK cells from COVID-19 participants (Figure 5A). We found 5 biologically relevant clusters among NK cells, of which C0 was significantly enriched in narrow neutralizers and C1 was significantly enriched in broad neutralizers (Figure 5B). NK cells in C0 are CD16^+^CD56^dim^ NK cells which express high transcript levels of *KLRK1* (NKG2D), *NCR1* (NKp46), *FASLG* (Fas-L), *LAMP1* (CD107a), *PRF1* (Perforin), and *GZMB* (Granzyme B), indicating cytotoxic potential. They also expressed intermediate levels of *CD69* and high levels of *CD38*, demonstrating activation. The C1 cluster (enriched in broad neutralizers) are also CD16^+^CD56^dim^, but exhibit lower levels of activation markers *CD38* and *CD69* as well as low expression of cytotoxic markers *KLRK1*, *NCR1*, *LAMP1*, and *PRF1*. However, the C1 cluster still highly expresses some cytotoxicity-related markers (*FASLG* and *GZMB)* and the cytokine *IFNG* (IFNγ) (Figure 5C). Thus, the cluster of NK cells enriched in broad neutralizers represents NK cells with diminished activation and more potential for cytokine secretion. The C1 cluster also expressed high levels of *MKI67* (Ki-67), further validating the increased proliferation of NK cells in broad neutralizers as compared to narrow neutralizers. The remaining clusters, C2-C4, are not differentially expressed between broad and narrow neutralizers.

**Figure 5:**
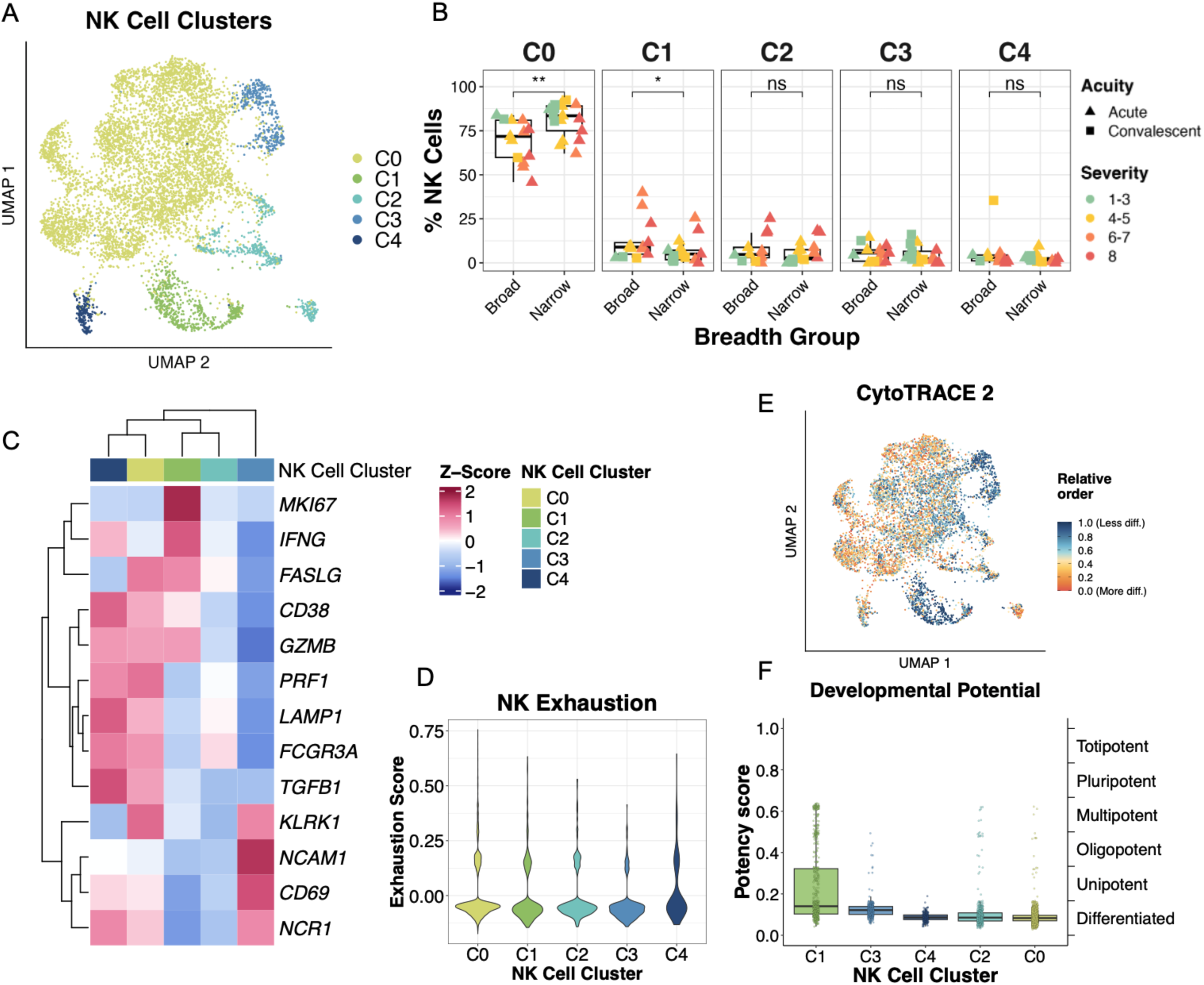
NK Cell Phenotype in Broad and Narrow Neutralizers. **A.** UMAP projection of NK cells colored by cluster. **B.** Boxplot of frequency of NK cells in each cluster split by breadth group. **C.** Heatmap of Z-score normalized gene expression of NK cell phenotypic and functional markers for each NK cell cluster. **D.** Violin plot of NK cell exhaustion (average expression of *LAG3*, *PDCD1*, and *HAVCR2*; see Materials and Methods) for each cell split by cluster**. E**. UMAP projection of NK cells colored by relative potency score calculated by CytoTRACE2. **F.** Boxplot of potency score of each NK cell cluster calculated by CytoTRACE2. *, P < 0.05; **, P < 0.01 by two-sided Wilcoxon rank-sum test with Bonferroni’s correction for multiple hypothesis testing.

In the setting of severe COVID-19, NK cells can become functionally exhausted while still expressing cytotoxic molecules^45–49^. Thus, we evaluated whether the highly activated cells in narrow neutralizers might be exhausted. We analyzed aggregated expression of canonical exhaustion markers (*LAG3*, *PDCD1* and *HAVCR2*) to generate an exhaustion score (Figure 5D). Although all clusters exhibited some evidence of exhausted cells, the C0 cluster enriched in narrow neutralizers was not predominantly composed of cells expressing exhaustion proteins. Thus, the expression of cytotoxic proteins observed in C0 is distinct from the exhaustion signal observed in severe COVID-19, in which NK cells do not retain intact cytotoxic function. Finally, we applied CytoTRACE2, a predictive method to evaluate absolute developmental potential in scRNA-seq data, to all COVID-19 NK cells^50^ (Figure 5E). We found that C1 contained cells with the highest developmental potential in this model, while C0 was the most differentiated, further supporting the observation that broad neutralizers are associated with immature NK cells and narrow neutralizers with differentiated NK cells (Figure 5F).

### Cell-cell interaction predictions indicate differences in cell signaling between broad and narrow neutralizers

Given that multiple immune cell types in narrow neutralizers demonstrated inflammatory phenotypes, we next investigated how cell-cell interactions may contribute to differential NK cell phenotypes in broad and narrow neutralizers. Our lab has previously found that monocytes contribute to NK dysfunction in severe COVID-19 by interacting with NK cells both through direct receptor-ligand interaction and via cytokine secretion^51^. We found that monocytes in narrow neutralizers express significantly higher levels of LLT-1 compared to broad neutralizers (Figure 6A). LLT-1 can activate NK cells and is downregulated in moderate and severe COVID-19^27^. LLT-1 binds CD161 on NK cells which has increased median expression in narrow neutralizers, but this difference is not significant (Figure 6B). ULBPs 1-2-5-6, ligands for the NK cell activating receptor NKG2D, were elevated in monocytes from broad neutralizers, while CD112 and CD155 (ligands for DNAM-1, TIGIT, and/or CD96 on NK cells) were not differentially expressed between breadth groups (Figure 6A). Cognate NK cell receptors NKG2D, DNAM-1, TIGIT, and CD96 were not differentially expressed between breadth groups on NK cells (Figure 6B).

**Figure 6:**
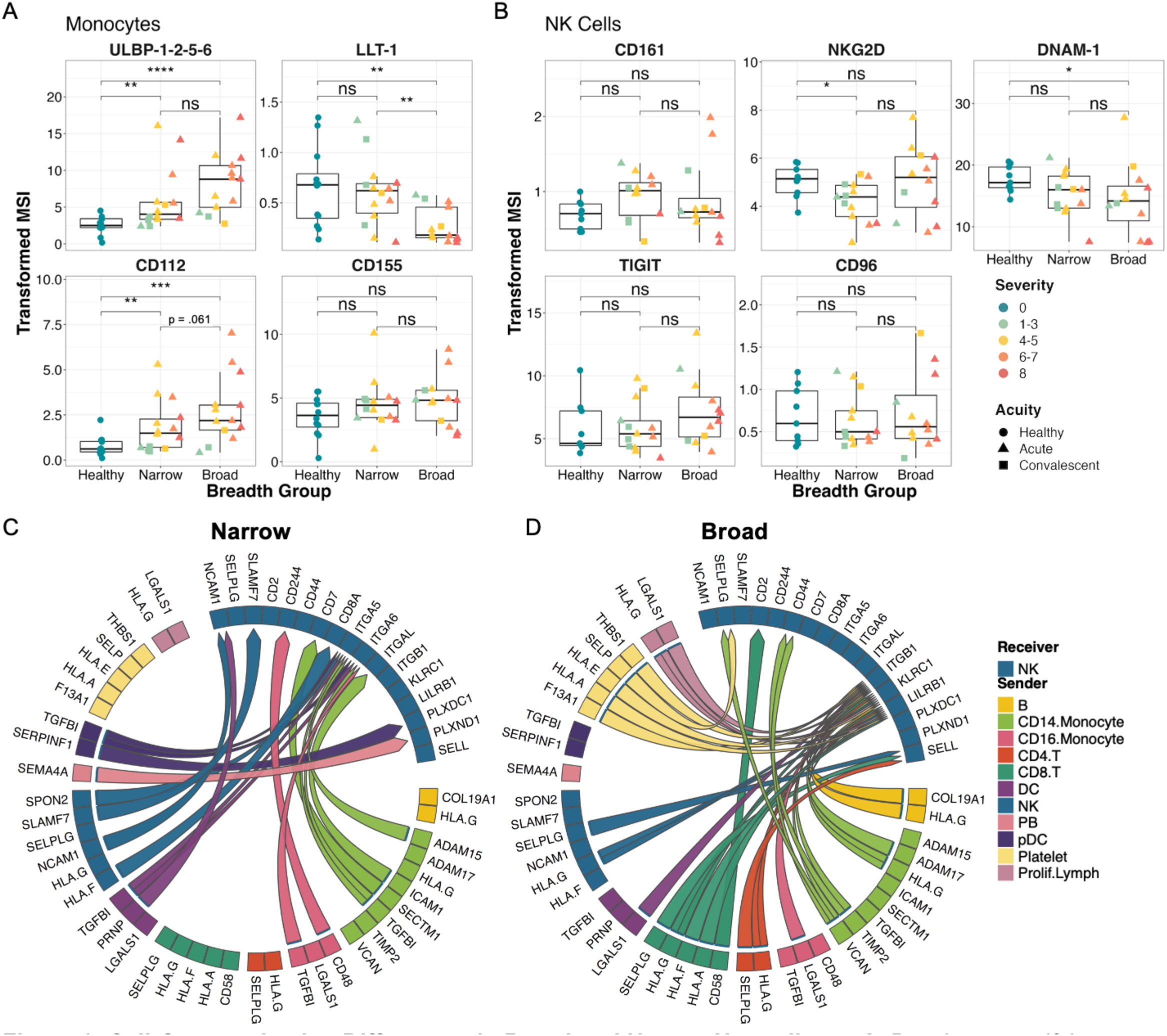
Cell-Communication Differences in Broad and Narrow Neutralizers. **A**. Boxplots quantifying arcsinh-transformed average expression of NK Cell Ligands on Monocytes in CyTOF data. **B**. Boxplots quantifying arcsinh-transformed average expression of cognate NK Cell receptors from CyTOF dataset. **C-D.** Circos plot showing top 50 predicted cell-cell communication pairs sent NK cells colored by sender cell in narrow (C) and broad (D) neutralizers.

To predict cell-cell communication from scRNA-seq data, we used the Multinichenetr package to identify the top differentially expressed receptor-ligand interaction pairs between breadth groups. The Multinichenetr method leverages the downstream signaling database Nichenet-v2 which integrates ligand-receptor, signaling, and gene regulatory data into a network model which allows linkage of ligand-receptor interactions to active downstream signaling^52^. Cell type and donor specific expression of ligand, receptor, and predicted downstream genes are used to rank interactions.This allows us to predict receptor-ligand interactions likely to be actively signaling in scRNA-seq data and thus could explain differential NK cell phenotypes between breadth groups (Figure 6C-D).

There were distinct patterns of predicted cell-cell communication received by NK cells in broad and narrow neutralizers. In narrow neutralizers, multiple cell types (CD14 monocytes, CD16 monocytes, pDCs, platelets, and DCs) were predicted to signal via *TGFBI* (TGFβ induced protein) through integrins (*ITGA5* and *ITGA6*) which inhibits adhesion and migration. We also investigated predicted downstream signaling targets of both TGFβI and TGFβ in the top 50 predicted cell-cell communication pairs in narrow neutralizers. TGFβI (*TGFBI*) had low regulator potential to drive ISG expression, but TGFβ (*TGFB*) had one of the highest predicted regulatory potential and is predicted to drive expression of many ISGs including *MX1* and *ISG15* (Extended Data Figure 4A). It is known that TGFβ in the context of viral infection and in conjunction with *IRF7* expression can drive type I interferon and ISG expression^53,54^. Here, *IRF7* was significantly upregulated in narrow neutralizers which may work in conjunction with TGFβ signaling to drive interferon-stimulated gene expression in narrow neutralizers. Other sources of activation included *CD48* interaction with 2B4 (*CD244)*^55^ on CD16 monocytes and *CD7* mediated activation by CD14 monocytes through *SECTM1*^56^. NK cells themselves were also predicted to send auto-activation signals via *SLAMF7*-SLAMF7^57,58^, CD56-CD56 (*NCAM1*)^59^, and *CD8*-*HLA-F*^60^ interactions. There were also predicted interactions between *SPON2* and Integrin Alpha 5 (*ITGA5*) in NK cells from narrow neutralizers, which can inhibit migration^61^. Additionally, signals involved in inhibition of angiogenesis (*SEMA4A*-*PLXND1*^62^ and *SERPINF1*-*PLXDC1*^63^) were predicted from pDCs and plasmablasts (Figure 6C).

CD4 T cells, CD8 T cells, proliferating lymphocytes, platelets, and B cells only were predicted to communicate with NK cells in broad neutralizers. Here, signaling was dominated by inhibitory receptors and HLA-E, HLA-A, HLA-F, and HLA-G were predicted to signal in multiple cell types through NKG2A (*KLRC1*) and/or LILRB1^37,41,64–68^. Receptor-ligand pairs involved in adhesion and lymphocyte homing were also predicted with signals from Galectin 1 (*LGALS1*) from proliferating lymphocytes and CD16 monocytes, Selectin P Ligand (*SELPLG*) from NK cells, CD4 T cells and CD8 T cells, Versican (*VCAN*) from monocytes, Selectin P (*SELP*) from platelets, and other signals through integrins across multiple cell types^69–72^. Some of the integrin-mediated signaling predicted in broad neutralizers are known to be inhibitory, such as *TIMP2* interaction with Integrin Beta 1 (*ITGB1*)^73^. Platelets are predicted to send inhibitory signals in broad neutralizers via Selectin P (*SELP*), *HLA-A*, *HLA-E*, and *THBS1*. Additional relevant signaling included immune synapse proteins *CD58* and *CD2* and *TIMP2*-*CD44* which can also be involved in migration and activation^74,75^(Figure 6D). Overall, these findings suggest that signals sent to NK cells in narrow neutralizers drive activation, including ISG expression as well as inhibition of migration and adhesion, and broad neutralizers are characterized by dominant inhibitory signals as well as positive regulation of migration and immune synapse formation.

### IFNα activated NK cells exhibit enhanced killing and upregulate cytotoxic markers when co-cultured with induced T follicular helper-like cells

The activation of NK cells in peripheral blood in narrow neutralizers raises the hypothesis that NK cells may target Tfh in lymphoid tissue to limit antibody development. Therefore, to directly assess the functionality of NK cells with narrow neutralizer phenotype against Tfh, we developed an *in vitro* co-culture system with healthy human donor NK cells and induced T follicular helper-like cells. Bonafide Tfh cells cannot be differentiated *in vitro*, but activation of primary, naive CD4 T cells with TGFβ and IL-12 has been shown to induce high expression of CXCR5, ICOS, and *BCL6* which indicates recapitulation of key features of Tfh cells^76^. After isolation of naive CD4 T cells from cryopreserved, healthy PBMCs, cells were activated with Staphylococcal Enterotoxin B (SEB; a T cell superantigen) for 4 hrs, then TGFβ and IL-12 were added to the culture for 72 hours, and CXCR5+ cells were sorted from this population. This pure population of CXCR5+ CD4 T cells is referred to here as induced T follicular helper-like (iTfh-like) cells (Figure 7A). These iTfh-like cells were confirmed to express high levels of ICOS and PD-1 indicating a Tfh-like phenotype (Extended Data Figure 5D).

**Figure 7:**
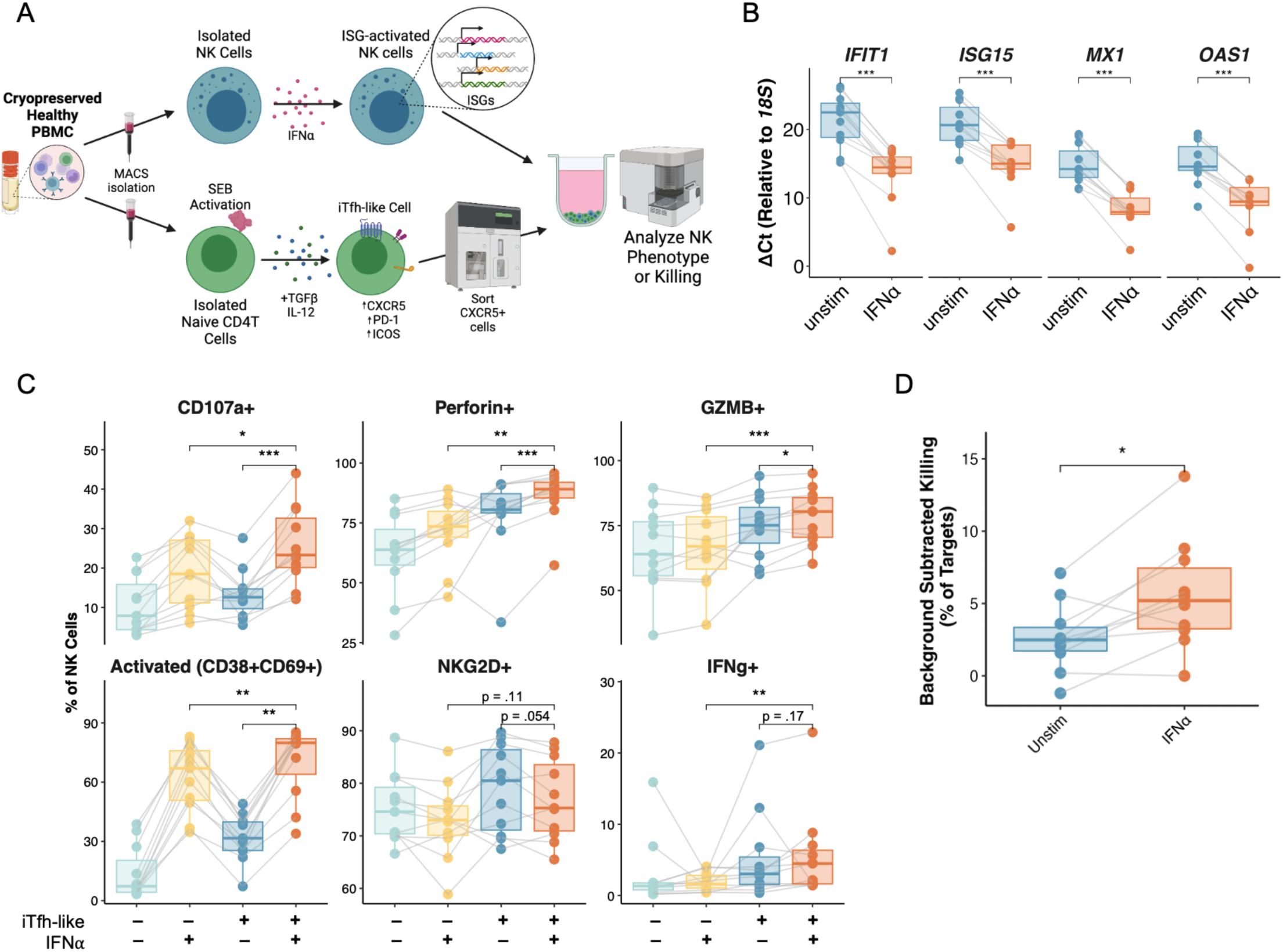
IFNα-Activated NK Cells Exhibit Increased Killing and Markers of Degranulation against iTfh-like Cells. **A.** Experimental design of in-vitro co-culture and killing experiments with healthy IFNα activated NK cells cultured with iTfh-like cells. **B.** Boxplots of ΔCt relative to *18S* of *IFIT1*, *ISG15*, *MX1*, and *OAS1* in healthy donor NK cells after activation with IFNα or unstimulated for 24 hrs. Each data point represents the average of 3 replicates from each donor. Lines connect donors. **C.** Quantification of percent of NK cells expressing cytotoxic and activation markers upon 16 hr culture with and without iTfh-like cells with and without NK cell pre-activation with IFNα for 24 hrs prior to co-culture at E:T of 1:5. Data shown are from n = 11 healthy donors across 3 separate experiments with lines connecting donors. **D.** Background-subtracted percent of iTfh-like cell death measured by ViaDye Red after 3 hr killing assay with either IFNα activated or unstimulated NK cells at an E:T ratio of 5:1. Cell death in targets-only well was used as background cell death. Data shown are from n = 10 unique healthy donors across 3 separate experiments with lines connecting unique donors. *, P < 0.05; **, P < 0.01; ***, P < 0.001 by two-sided, paired Wilcoxon rank-sum test with Bonferroni’s correction for multiple hypothesis testing.

In order to mimic the high ISG expression in narrow neutralizer NK cells, we activated NK cells with IFNα to induce ISG expression. *In vitro* treatment of NK cells with IFNα has been reported to recapitulate ISG expression observed in COVID-19 participants^46^. We found that in isolated healthy donor NK cells, 24 hours of activation with IFNα led to significantly greater expression (lower ΔCt relative to *18S*) of hallmark ISGs: *MX1*, *OAS1*, *ISG15*, and *IFIT1* (5-8 ΔCt) measured by RT-qPCR (Figure 7B). We then investigated how NK cells pre-activated with IFNα responded to iTfh-like cells by co-culturing them at an Effector:Target (E:T) ratio of 1:5 overnight (Figure 7A). Following co-culture, significantly more NK cells pre-activated with IFNα expressed CD107a (median % positive: unstimulated = 13.0%, IFNα = 23.3%), Perforin (unstimulated = 80.4%, IFNα = 89.1%), and Granzyme B (unstimulated = 75.3%, IFNα = 80.4%) compared to unstimulated NK cells in co-culture and IFNα pre-activated NK cells incubated without target cells (Figure 7C). This trend was also observed in the percent of activated (CD38, CD69 double positive) NK cells (unstimulated = 35.0%, IFNα = 79.9%). IFNα pre-activated NK cells in co-culture also expressed more IFNγ than IFNα pre-activated cells cultured alone but this difference was not significant between IFNα pre-activated and unstimulated NK cells in co-culture. The proportion of NKG2D+ NK cells was slightly decreased in IFNα pre-activated co-cultured NK cells, although this difference was not significant^77^ (Fig 7C).

We also performed killing assays with IFNα pre-activated NK cells against iTfh-like cells at an E:T ratio of 5:1 for 3 hours. After incubation, a flow-based, fixable viability dye was used to evaluate iTfh-like cell death after co-culture with IFNα pre-activated NK cells or unstimulated NK cells as well as background cell death in an iTfh-like only condition. After subtracting background cell death in target-only wells, we found that IFNα pre-activated NK cells killed significantly more iTfh-like cells than unstimulated NK cells (median difference 2.7%) (Fig 7D). We further determined that this killing was iTfh-like specific because NK cells activated with IFNα did not kill significantly more magnetically-isolated bulk blood CD4 T cells (Extended Data Figure 5C). Finally, we found that the magnitude of upregulation of ISGs for each healthy donor measured by ΔΔCt from RT-qPCR (the absolute value of the difference in ΔCt values between IFNα activated and unstimulated NK cells) correlated with the difference in killing of iTfh-like cells between IFNα pre-activated and unstimulated NK cells (ΔKilling), with greater cytotoxicity associated with greater ISG gene expression. This correlation was only statistically significant for *OAS1* (R = 0.69, P = 0.028) and other genes approached significance (Extended Data Figure 5E). Overall these data suggest that NK cells with narrow neutralizer phenotype have capacity to specifically target autologous Tfh, potentially driven by ISG expression.

## Discussion

The COVID-19 pandemic has highlighted the critical need to understand immunological determinants of antibody breadth to formulate actionable strategies to elicit broad protection with vaccines. Although NK cell activation has been repeatedly highlighted as a correlate of poor response to vaccines^8,9,16^, there is a dearth of research to elucidate the mechanism of human NK cell regulation of Tfh and antibody responses, particularly in the context of acute infection. We leveraged multi-omic approaches and our unique cohort of individuals experiencing primary infection with ancestral SARS-CoV-2 in combination with *in vitro* functional validation to uncover that type I interferon activation and greater proportion of NK cells is correlated with narrow antibody response, whereas features of immaturity and inhibitory signaling in NK cells is correlated with broad antibody responses. We further demonstrate that interferon-driven activation of NK cells increases their response to and killing of autologous iTfh-like cells, our hypothesized mechanism of NK cell regulation of antibody breadth.

Interferon-stimulated genes (ISGs) are expressed in response to interferons such as interferon alpha and gamma and have critical antiviral functions^78^. In COVID-19, interferon signaling and its role in pathology or viral clearance has remained controversial, with early interferon signaling thought to promote viral clearance and persistent interferon signaling leading to greater disease severity^79–81^. Beyond its association with severity, we found an alternate role for interferon-mediated activation in NK cells: we showed, through multiple overlapping methods, that NK cells in narrow neutralizers show profound type I interferon activation. The abundant NK cells from narrow neutralizers exhibited significant upregulation of many ISGs including the IFI family, MX family, OAS family, *ISG15*, etc. In fact, 81/86 genes that were upregulated in narrow neutralizers were ISGs according to the interferome database^30^. NK cell clusters enriched in narrow neutralizers expressed multiple cytotoxic proteins and were the most differentiated of the NK cell clusters in our dataset. These NK cells also had migratory potential to traffick to lymphoid tissues due to their expression of CX3CR1 which is a chemokine receptor that also defines highly cytotoxic, mature NK cells^31^. Overall, our data links ISG-mediated activation with increased cytotoxic potential of NK cells against Tfh. We then validated this *in vitro* by demonstrating that ISG-activated NK cells exhibit increased cytotoxic activity against autologous iTfh-like cells. While the difference in killing by ISG-activated and resting NK cells is relatively small (median difference = 2.7%), this is consistent with a level of killing that could drastically affect antibody responses where few Tfh are responsible for amplifying antigen-specific antibody responses. Tfh are the limiting factor in germinal center reactions and intervention at this point in the immune response is greatly amplified over the course of viral infection^82^. The level of iTfh-like killing in IFNα-activated NK cells was correlated with ISG upregulation, supporting a proposed mechanism where ISG-mediated activation in NK cells leads to killing of autologous cells. Additionally, killing of iTfh-like cells appeared specific with no difference in killing against bulk blood CD4 T cells.

In broad neutralizers, NK cells exhibited evidence of immaturity and proliferation with no ISG activation. NK cells in broad neutralizers expressed higher levels of CD94, NKG2A, and CD127 (*IL7R*) in CyTOF and/or scRNA-seq data which can indicate both immaturity as well as inhibitory signaling. This inhibitory signaling was doubly revealed through our predicted cell-cell communication analysis with active signaling via *KLRC1* (NKG2A) and *LILRB1*. LILRB1 engagement can even directly decrease granzyme B and cytotoxicity, which could prevent NK cells from restricting antibody responses^83,84^. NKG2A and CD127 are also features of immature NK cells which was reinforced by analysis of developmental potential using CytoTRACE2 and pathway analysis. NKG2A, which dimerizes with CD94 to send inhibitory signals, has been identified to play a role in multiple aspects of the productive immune response to SARS-CoV-2. SARS-CoV-2 viral peptides stabilize HLA-E (NKG2A/CD94 ligand) and prevent binding to NKG2A which “unleashes” NKG2A+ NK cells to eliminate SARS-CoV-2 infected target cells^85^. So even as it is NKG2A could be restricting NK cell killing of Tfh or regulation of adaptive immunity, these NKG2A+ cells could be unrestricted in killing of virally infected cells, leading to clearance of the virus. Furthermore, NKG2A was also identified as a predictor of response to the Moderna mRNA vaccine against SARS-CoV-2, indicating the clinical relevance of this trend^86^. HLA-Bw4, but not HLA-Bw6 expression levels were also elevated in broad neutralizers. HLA-Bw4 has been associated with resistance to AIDS and clearance of SARS-CoV-2^42,43^. We further suggest HLA-Bw4 may be associated with more productive antibody responses.

Our study represents the first association between NK cell responses and antibody breadth in acute human viral infection. Though we show that ISG-expression in NK cells drives killing and response to iTfh-like cells further work is needed to identify NK cells in lymphoid tissues after vaccination and infection to confirm this model. We were also unable to sequence the B cell receptor repertoire in our cohort due to sample availability. Sequencing and epitope mapping are needed to precisely describe the NK cell mediated effect on antibody breadth and to determine its effect on somatic hypermutation directly.

Here, we used a systems immunology approach to identify a potential interferon-mediated mechanism driving NK cell activation and killing of autologous Tfh cells which may be the link between NK cells, poor vaccine response, and limited antibody breadth. We additionally identify features of immaturity and inhibitory signaling in broad neutralizers of SARS-CoV-2, including NKG2A. Understanding these positive and negative correlates of breadth are critical to developing durable, broad-spectrum vaccines against COVID-19 and other emerging pathogens. COVID-19 vaccines were developed and entered clinical trials only 128 days after the COVID-19 global pandemic was declared, but by the time Omicron B.1.1.529 emerged, efficacy estimates dropped as low as 9% and the virus had evaded most antibody therapies available^87–89^. This underscores the need to understand broad antibody responses against novel variants so that we can formulate actionable strategies to elicit broad protection with vaccines. In the future, we can use this understanding of the acute immune response against SARS-CoV-2 that leads to broad and narrow neutralization to select and designs adjuvants and immunogen combinations that minimize ISG-activation and maximize the potential of NKG2A+ NK cells both against virally infected cells and in driving favorable antibody responses.

## Methods

### Pseudotyped Virus Production

SARS-CoV-2 variant spike pseudotyped lentiviral particles were produced in 293T cells (ATCC) using Fugene transfection reagent 6 (Promega; E2691). Two million cells were seeded in D10 (DMEM(Life Technologies; 11885-092) + 10% FBS, 1% L-glutamine, 1% PSA, 10mM HEPES) in T75 flasks 24 hrs before transfection. A 3-plasmid system was used for viral production, as described Ou et al. 2020^90,91^. The spike vector contained the 21 amino acid truncated form of the SARS-CoV-2 sequence from the Wuhan-1 strain of SARS-COV2. Wu-1 Sequence ID: BCN86353.1; Alpha – sequence ID: QXN08428.1; Beta – sequence ID: QUT64557.1; Delta – sequence ID: QWS06686.1, which also has V70F and A222V mutations; and Omicron – sequence ID: UFO69279.1. In a final volume of 1mL, 1μg Lenti backbone, 1.28 μg psPAX2, and .34μg of spike plasmid were mixed with serum-free DMEM. 100μl of fugene6 was added dropwise and incubated at room temperature for 15 min to form transfection complexes. The 1mL transfection mix was then added dropwise to plated 293Ts. Culture medium was removed and replaced 24hrs after transfection. Viral supernatants were harvested 72 hrs after transfection by filtering through a .45μM filter. Viral stocks were aliquoted and stored at −80C until further use (no less than 48hrs).

### Pseudovirus Neutralization Assay and Breadth Score

Target cells were from HeLa Cell line stably overexpressing ACE2 and TMPRSS2 produced as described in Rogers et al. 2020^90^. HeLa-ACE2-TMPRSS2 cells were thawed in D10 and passaged at least twice before use. 96-well black walled, clear bottom plates were used for the assay (Thermo Fisher; 165305). Patient serum was inactivated at 56C for 1hr before assay was performed. Patient serum was diluted 1:50 in D10 (final dilution of 1:100 when mixed with virus) and filtered through a .22uM spinX filter (Corning; 8160) and further serial dilutions (1:2) up to 1:1600 (final dilution 1:3200) were performed on the plate in 25μL final volume. Diluted patient sera was mixed with 25μL of pseudovirus at a titer such that virus-only wells would achieve a luminescence of approximately 100,000 RLU on Promega Glomax plate reader (Promega; GM3000). Virus-serum mixtures were incubated for 1hr at 37C. During incubation, HeLa-ACE2-TMPRSS2 cells were harvested using trypLE (Gibco; 12604021) and resuspended at .2e6 cells/mL. After incubation 50μL cells were added to each well (10,000 cells per well). After 48 hrs of incubation, all media was removed from plate by vacuum aspiration and cells were lysed by addition of 50μL of britelite plus assay readout solution (Revvity; 6066766) and 50μL of 1X PBS. Luminescence was measured using Promega Glomax after 30s shaking. Background luminescence (cell only wells) were averaged and subtracted from all wells. Percent neutralization was calculated by comparing to 100% infection (virus only wells).The reciprocal of the highest dilution to obtain <50% infection was used to calculate average half maximal neutralization titer (NT50). The average NT50 for each patient against each virus was calculated using 3 technical replicates per virus. Number of average NT50s above 100 (limit of detection) were used as Breadth Score.

### Processing of scRNA-seq

PBMC and whole blood Samples were collected and processed in seqwell-based scRNA-seq as described in Wilk et al. 2021. From publicly available data, the R package Seurat (V4.4.0)^40^ was used to subset participants included in this study and to remove neutrophils and developing neutrophils as they were not preset for each participant. Subsetted data was re-scaled and transformed using SCTransform() function, and linear regression was performed to remove unwanted variation due to cell quality (e.g., percentage mitochondrial reads, percentage rRNA reads, percentage hemoglobin genes in whole blood). Principal component (PC) analysis (PCA) was performed using the 3,000 most highly variable genes, and the first 50 PCs were used to perform Uniform Manifold Approximation and Projection for Dimension Reduction (UMAP) to embed the dataset into 2 dimensions. Next, the first 50 PCs were used to construct a shared nearest neighbor graph (SNN; FindNeighbors()). Cell types identified in Wilk et al. were verified using known lineage markers. Differentially expressed genes for each cell type were identified using Seurat’s FindMarkers() using wilcoxon rank sum test, log_2_FC > .25 and adjusted P-value < .05 (Table 3).

### DESeq2 Correlation Analysis

Raw counts from Seurat objects were subsetted by removing cells with less than 300 reads, more than 4000 unique reads, more than 10000 total reads, greater than 15% mitochondrial reads, less than 1% mitochondrial reads, greater than 7% ribosomal reads, and greater than 50% rrna. Raw pseudobulk count matrices were constructed using the AggregateExpression() function in Seurat to create counts per donor. Using DESeq2 (V1.38.3), these count matrices (from whole PBMCs or annotated NK cells) were used to make a DESeq object (DESeqDataSetFromMatrix()) with design = ∼Score and differentially expressed genes that correlate with breadth score as a numerical variable were identified using DEseq() with adjusted P-value < .05^29^.

### KEGG Pathway analysis

Stringdb package (V2.10.1) was used to calculate gene set enrichment of differentially expressed genes from the KEGG pathway database^44,92^. All genes measured in scRNA-seq were used to set background using stringdb$set_background(). Upregulated genes in each breadth group were mapped to STRING IDs using string_db$map() and enrichment was calculated using string_db$get_enrichment() hypergeometric test with Benjamini-Hochberg multiple hypothesis testing correction.

### Gene Module Scoring

The Seurat function AddModuleScore() was used to score single cells by expression of a list of genes of interest. This function calculates a module score by comparing the expression level of an individual query gene to other randomly selected control genes expressed at similar levels to the query genes and is therefore robust to scoring modules containing both lowly and highly expressed genes, as well as to scoring cells with different sequencing depths.

### Bionet

A protein-protein interaction network of the 2000 most differentially expressed genes in NK cells was used as the background interaction graph constructed using STRINGdb. Genes were identified from a Seurat object on NK cells from only COVID-19 participants using FindVariableFeatures() with selection.method = vst and n.features = 2000. Our STRINGdb background graph is available in our github repo (see Data Availability). The Bionet R package (V1.58.0) to find the highest-scoring subgraph of differentially expressed genes^33,34^. Nodes are scored using a β-uniform mixture model fitted to the adjusted P-value distribution (fitBumModel(), scoreNodes(FDR = 1e-6)). The minimal significant network was calculated using runFastHeiz() and nodes were colored by logFC from differential expression analysis.

### MultiNicheNet Analysis

MulitnichenetR (V1.0.1)^52^ was used to identify differentially expressed active receptor-ligand interactions between broad and narrow neutralizers. Top 50 receptor-ligand interactions received by NK cells were identified using default parameters (logFC_threashold = .5, p_val_threashold = .05, fraction_cutoff = .05). These receptor-ligand interactions were manually verified by searching the literature for publications establishing receptor-ligand interactions between two cells.

### Cell Developmental Potential Score (cytoTRACE2)

CytoTRACE2 (V1.0.0) classifies cells’ developmental potential using an interpretable deep learning framework. Default parameters were used to run to score each NK cell using cytotrace2() function on RNA counts using the “human” model^50^.

### NK Isolation and Activation

NK cells were isolated from cryopreserved, healthy PBMCs. PBMCs were thawed at 37 C and washed with complete RP10 (RPMI (Gibco;11875093), 10% FBS, 1% PSA, 1% L-glutamine, 10mM HEPES, 1% NEAA, 1% Sodium Pyruvate). Untouched, magnetically isolated NK cells were isolated using Miltenyi NK Cell Isolation Kit, human (Miltenyi; 130-092-657) according to manufacturer’s protocol. After isolation, NK cells were incubated for 24hr with 5000U/mL rh-IFNα A/D (pbl; 11200-1) in complete RPMI in 96 well U-bottom plates.

### Induced T Follicular Helper-Like Cell Differentiation

Induced T follicular helper-like (iTfh-like) cells were generated based on Schmitt et al. 2014 with some differences^76^. Naive CD4+ T cells were isolated from cryopreserved PBMCs using Miltenyi Naive CD4+ T Cell Isolation Kit II, human (Miltenyi; 130-094-131) using 1.1x manufacturers recommendation for antibody cocktail and biotin beads. Isolated cells were resuspended at 2e6 cells/mL in complete RPMI and activated with 1μg/mL Staphylococcus Enterotoxin B (Toxin Technologies; NC9442400) for 4 hrs at 37C. TGFβ (R&D Systems; 7754-BH-005, 5ng/mL) and IL-12 (R&D Systems; 219-IL-005, 1ng/mL) were added to the culture for 72 hrs. Cells were harvested and stained for CXCR5 expression for 15 minutes at 37C by adding 30μL of anti-human CXCR5 AlexaFluor 488 (Biolegend; 356912) to dry pellet. 1mL of complete RPMI was added and cells were spun down and resuspended in complete RPMI at 5e6 cells/mL. CXCR5+ cells (iTfh-like cells) were sorted using Sony SH800 into complete RPMI with 50% FBS. Cells were counted to ensure viability. Cells before and after sorting were saved for staining of Tfh markers. Unlabeled iTfh-like cells were plated before and after sorting and cells were washed in FACS (1XPBS +.05% BSA+2% FBS) twice and resuspended in ViaDye Red (Cytek; R7-60008) at 1:6000 dilution in PBS. After 30 min at room temp, cells were washed twice with FACS and resuspended in a cocktail of surface antibodies (anti-CXCR5 Alexa Fluor 488, anti-PD-1/CD279 BV711, anti-ICOS/CD278 BV711, anti-CD4 cfluor V610) for another 30 min at room temperature. After surface stain, cells were washed twice in FACS and fixed in 2% paraformaldehyde (EMS; 15710) for 10 minutes at RT. Cells were then resuspended in FACS and analyzed on Cytek Aurora flow cytometer.

### NK Cell Functional Co-Culture and Phenotyping

After activation of NK cells and sorting of iTfh-like cells, iTfh-like cells were labeled with CellTrace Violet (Invitrogen; C34557) for 20 minutes at 37 C at 1:10,000 dilution in PBS. After labeling, 5X volume complete RPMI was added for 5 min to remove any free dye. Cells were then pelleted, washed once with RPMI and resuspended for co-culture. NK cells and iTfh-like cells were added at an E:T ratio of 1:5 in 96-well U-bottom plates. Co-cultures were centrifuged at 1000 RPM for 1 min to bring cells together and incubated overnight before analysis of NK cells by flow cytometry. 4 hrs before the end of co-culture Brefeldin A (eBioscience; 00-4506-51), Monensin (eBioscience;00-4501-51), and anti-CD107a PE was added to the culture, followed by centrifugation at 1000 RPM for 1 min. When co-culture was complete, cells were washed in FACS twice and resuspended in ViaDye Red at 1:6000 dilution in PBS. After 30 min at room temperature, cells were washed twice with FACS and resuspended in a cocktail of surface antibodies (anti-NKG2D BV650, anti-CD38 SuperBright 600, anti-CD3 BV785, anti-CD14 BV785, anti-CD19 BV785, anti-CD69 PE-Dazzle594, anti-CD56 PE-Cy7, anti-TRAIL/CD253 APC, Anti-CD16 Alexa Fluor 700) in 1X Brilliant Stain Buffer (BD; 563794) diluted with FACS for another 30 min at room temperature. After surface stain, cells were washed twice in FACS and fixed in 2% paraformaldehyde for 10 minutes at RT. Cells were then washed twice in 1X permeabilization buffer (eBioscience; 00-8333-56) and stained with intracellular antibodies (anti-Perforin BV510, anti-IFNg BV711, anti-Granzyme B PerCP-Cy5.5) in 1X Brilliant Stain Buffer and 1X permeabilization buffer diluted in water for 30 min at RT. Cells were then washed twice with permeabilization buffer, resuspended in FACS and analyzed on Cytek Aurora flow cytometer.

### Flow Cytometry Based NK Cell Killing Assay against iTfh-like cells

After activation of NK cells and sorting of Tfh-like phenotype of NK cells, iTfh-like cells were labeled with CellTrace Violet as described above. NK cells and iTfh-like cells were added at an E:T ratio of 5:1 in 96-well U-bottom plates. Cells were centrifuged at 1000 RPM for 1 min to bring cells together and incubated at 37 C for 3 hrs. Cells were then washed twice in FACS and resuspended in ViaDye Red at 1:6000 dilution in PBS. Cells were then washed twice in FACS and fixed in 2% paraformaldehyde for 10 minutes at RT, and resuspended in FACS for analysis on Cytek Aurora flow cytometer.

### Flow Cytometry Analysis

Spectral unmixing was performed in Cytek Spectroflo Software using UltraComp eBeads Plus Compensation Beads (Thermo Scientific; 01-3333-42) as single color controls. Unmixed FCS files were imported to FlowJo V10.10.0 and gated as described in Extended Data Figure 6 to identify cell populations and marker expression. All data shown also in Table 5.

### qPCR

After NK cells were isolated and activated with IFNα as described above for 24hrs, cells were lysed in DNA/RNA Shield (Zymo Research; R1100-250). RNA was isolated using RNA Clean and Concentrator and Concentrator kits (Zymo Research; R1018) and excess DNA was removed from the samples using the TURBO DNA-free Kit according to the manufacturer’s instructions (Fisher Scientific, Cat. AM1907). RT-qPCR reactions were prepared with 1ng RNA per well using the Invitrogen superscript III Platinum One Step qRT PCR Kit with ROX (Invitrogen; 11745500) and TaqMan Gene Expression assay FAM (Thermo Scientific, ISG15; 4331182, MX1; 4331182, OAS1; 4331182, IFIT1; 4331182) according to manufacturer’s protocol. QuantStudio 3 Real-Time PCR System was used to quantify transcript levels (Thermo Fisher; A28567). Three technical replicates of each sample were measured and all samples were normalized to an 18S endogenous control (Thermo Scientific; 4352930E). ΔCt was calculated by Ct (experimental gene) - Ct (18S). ΔΔCt was calculated by ΔCt(unstimulated NK cells) - ΔCt(IFNα-activated NK cells).

### Data Visualization

All data analysis and visualizations were performed in the open source software R (V4.2.2)^93^. The R package Seurat was used to generate UMAP projections. ComplexHeatmap (V2.14.0) was used for all heatmaps^94,95^. ComplexUpset (V4.2.2) was used for upset plot^96^. MultinichenetR was used to create circos plots. EnhancedVolcanoPlot (V1.16.0) was used for volcano plots. Colors for plots were generated using MoMAColors, MetBrewer, NatParkPalettes, and PNWColors. Custom ggplot2 functions were used for all other plots.

### Data availability

FCS files (mass cytometry) with de-identified metadata supporting this publication are available at ImmPort (https://www.immport.org) under study accession no. SDY1708. Data from scRNA-seq have been deposited with the Gene Expression Omnibus under accession no. GSE174072. FCS flow cytometry files from *in vitro* assays are deposited at CytoBank Community Experiment IDs: 118822, 118828, 118829, 118830. A Github repository for all code will be available upon publication at https://github.com/BlishLab/SARSCoV2_Antibody_Breadth.

## Extended Data

**Extended Data Figure 1:**
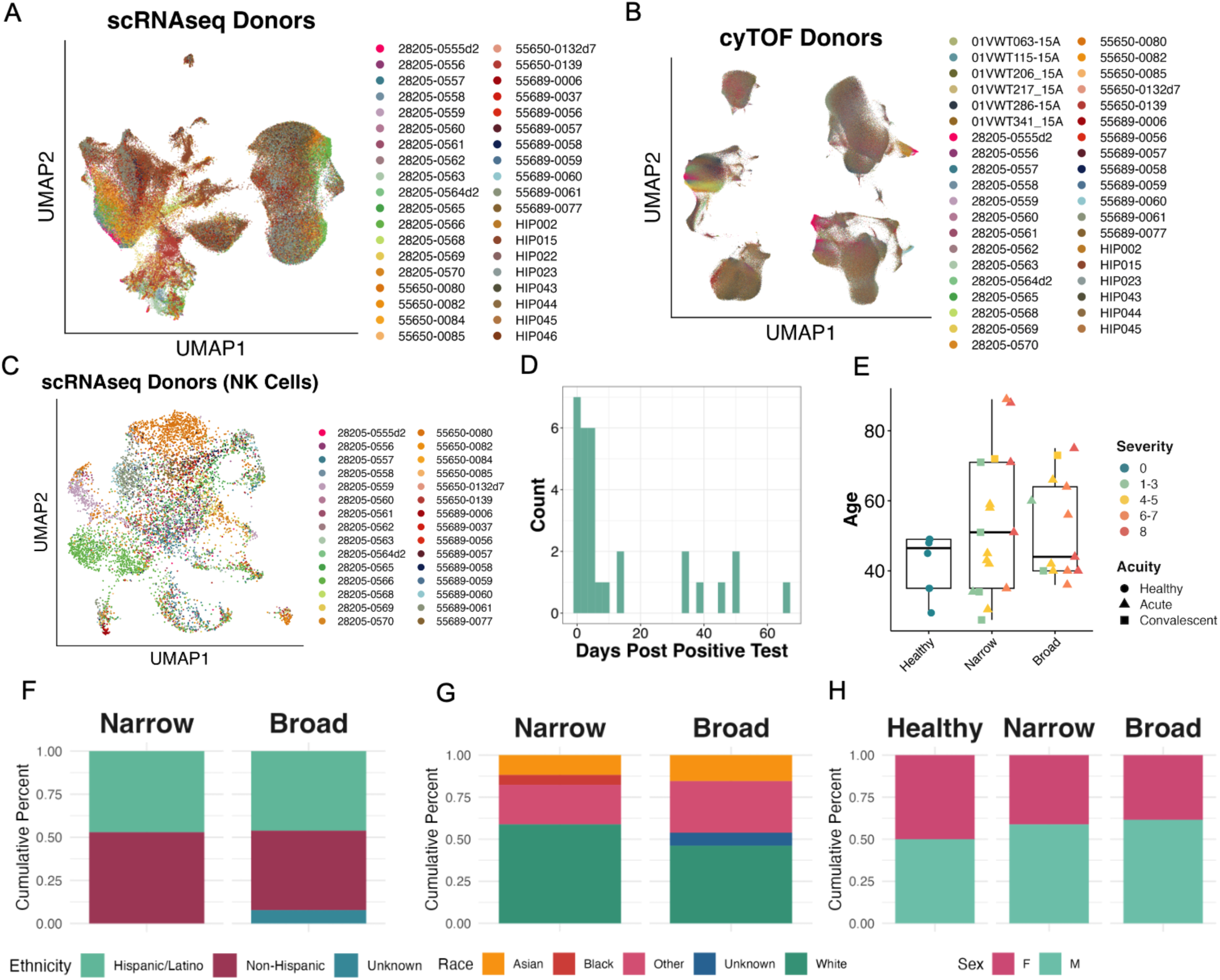
Cohort Characteristics. **A**. UMAP embeddings of complete scRNA-seq dataset, colored by donor. **B.** UMAP embedding of complete CyTOF dataset, colored by donor. **C.** UMAP embeddings of NK cells from all COVID-19 participants, colored by donor. **D.** Histogram of days post positive test for each sample. **E.** Box plot of age at blood draw for each donor in broad, narrow, and healthy controls. **F.** Cumulative bar plot of relative percent of reported ethnicity in broad and narrow breadth groups. **G.** Cumulative bar plot of relative percent of reported race in breadth groups. **H.** Cumulative bar plot of relative percent of reported sex in broad, narrow, and healthy groups. Ethnicity and race were not reported for healthy controls.

**Extended Data Figure 2:**
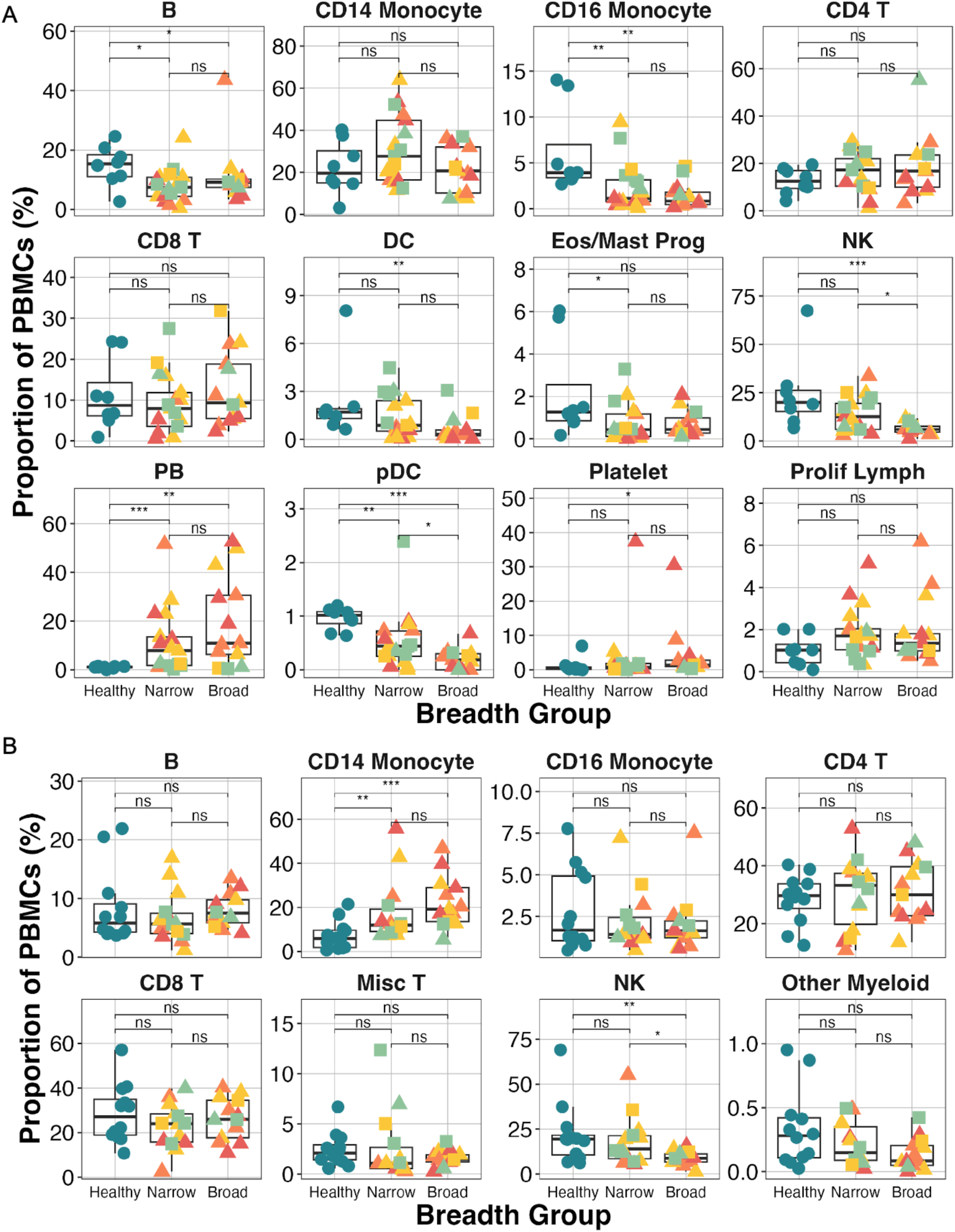
Cell Type Frequencies in Breadth Groups in scRNAseq and CyTOF datasets. **A.** Cell type proportions from scRNA-seq data of PBMCs in each sample. Platelets are excluded from the proportion calculations because their presence is related to sample processing. **B.** Cell type proportions from CyTOF data of PBMCs in each sample. P < 0.05; **, P < 0.01; ***, P < 0.001; ns, P = 0.05 by two-sided Wilcoxon rank-sum test with Bonferroni’s correction for multiple hypothesis testing. PB = plasmablast, Eos = eosinophil; Prog = progenitor; Prolif Lymph = proliferating lymphocyte.

**Extended Data Figure 3:**
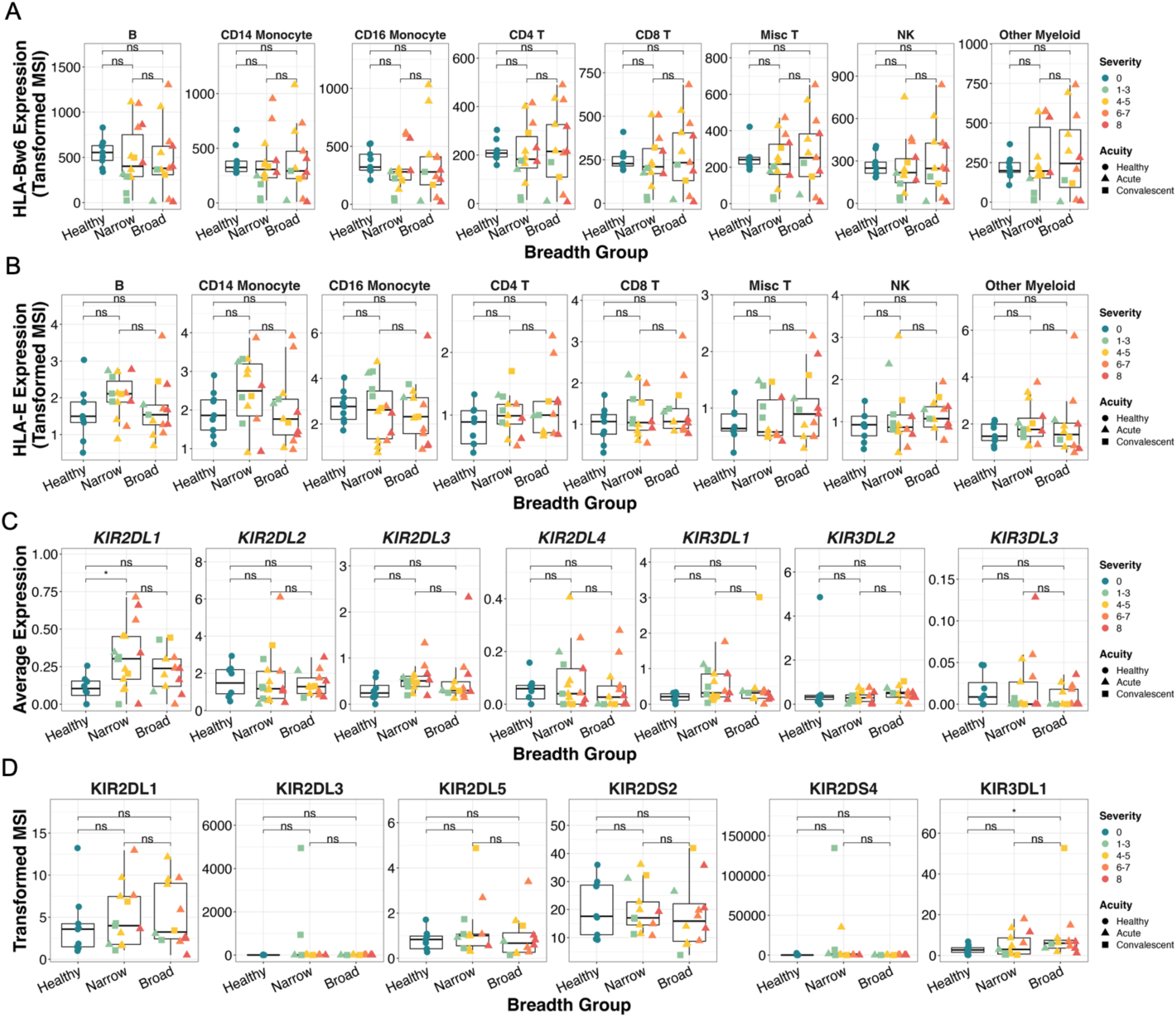
CyTOF and scRNA-seq Expression Data of HLA and KIR Genotypes. **A**. Boxplots quantifying arcsinh-transformed average expression of HLA-bw6 in each cell type from CyTOF dataset. **B**. Boxplots quantifying arcsinh-transformed average expression of HLA-E in each cell type from CyTOF dataset. **C**. Boxplots quantifying RNA expression of all KIR genes detected in scRNAseq dataset in NK cells. **D**. Boxplots quantifying arcsinh-transformed average expression of all KIR types included in CyTOF panel in NK cells. *, P < 0.05; **, P < 0.01; ***, P < 0.001 by two-sided Wilcoxon rank-sum test with Bonferroni’s correction for multiple hypothesis testing.

**Extended Data Figure 4:**
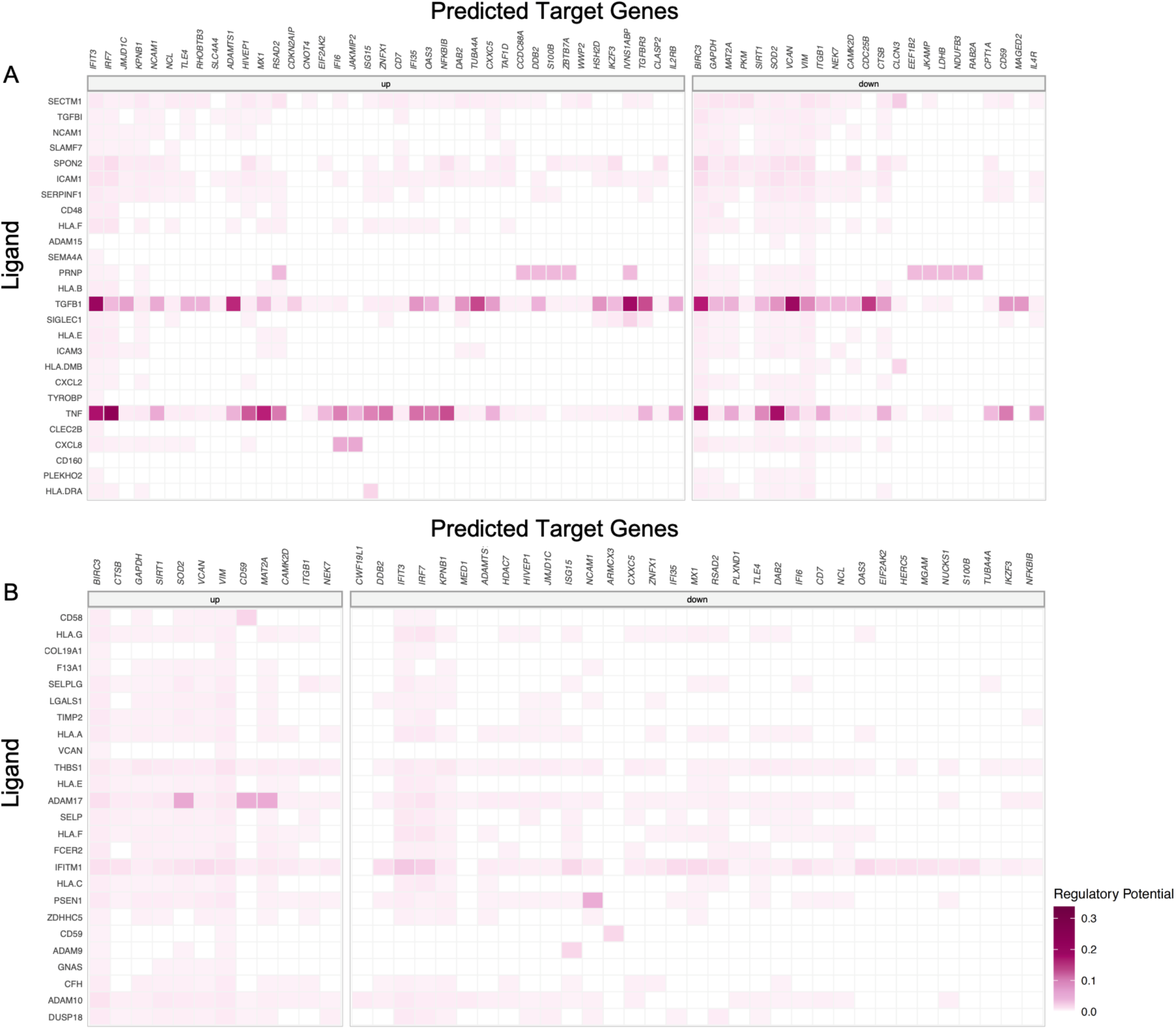
Predicted Target Gene Expression in Cell-Cell Communication Pairs in Narrow and Broad Neutralizers. **A and B.** Heatmap of scaled ligand activity, ligand activity, and regulatory potential of predicted downstream gene expression of the top 50 ligand-receptor pairs sent from all PBMC cell types to NK cells from narrow (A) and broad (B) breadth groups. Scaled ligand activity values are calculated as Z-score normalized NicheNet ligand activity values. Regulatory potential is derived from ligand-target regulatory priors model from nichenet-v2.

**Extended Data Figure 5:**
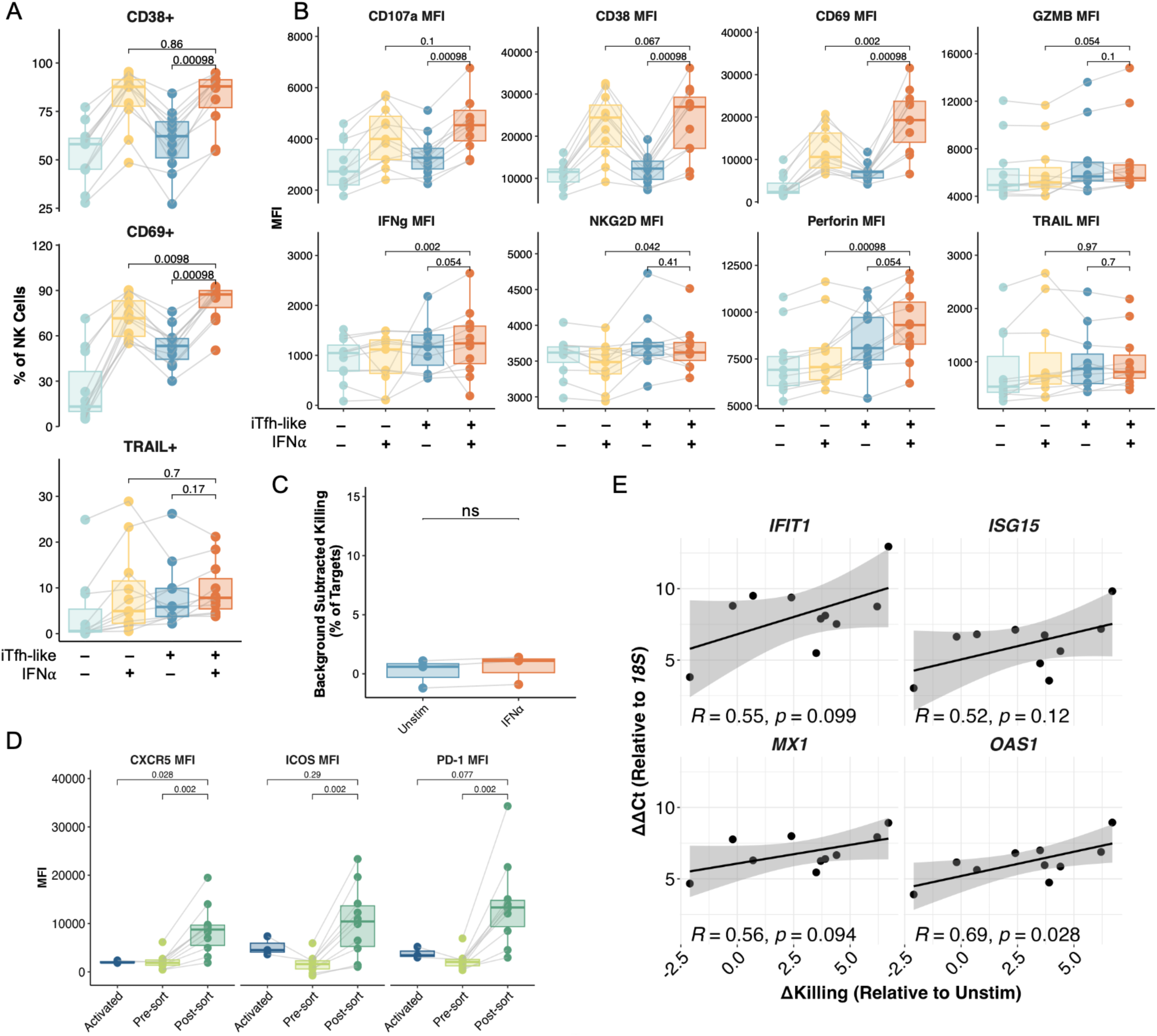
IFNα-Activated NK Cells Exhibit Increased Killing and Markers of Degranulation against iTfh-like Cells. **A.** Quantification of percent of NK cells expressing cytotoxic and activation markers upon co-culture with and without iTfh-like cells with and without NK pre-activation with IFNα for 24 hrs prior to co-culture at E:T of 1:5. **B.** Mean Fluorescence Intensity (MFI) of cytotoxic and activation markers in NK cells after co-culture with and without iTfh-like cells, with and without NK pre-activation with IFNα for 24 hrs prior to co-culture at E:T of 1:5. Data shown in A and B are from n = 11 healthy donors across 3 separate experiments with lines connecting donors. **C**. Background Subtracted percent of magnetically-isolated CD4 T cell death measured by viadye red with either IFNα activated or unstimulated NK cells with an E:T ratio of 5:1 **D**. Mean fluorescence intensity (MFI) of Tfh markers after activation (SEB, but no cytokines) and iTfh-like differentiation before and after sorting on CXCR5+ cells. Gated on live, CD4+ lymphocytes. **E.** Correlation between Δkilling (% killing IFNα pre-activated NK cells - % killing unstimulated NK cells) and ΔΔCt for expression of interferon-stimulated genes relative to unstimulated NK cells. P-values by Spearman rank correlation. Other P-values by two-sided, paired Wilcoxon rank-sum test with Bonferroni’s correction for multiple hypothesis testing.

**Extended Data Figure 6:**
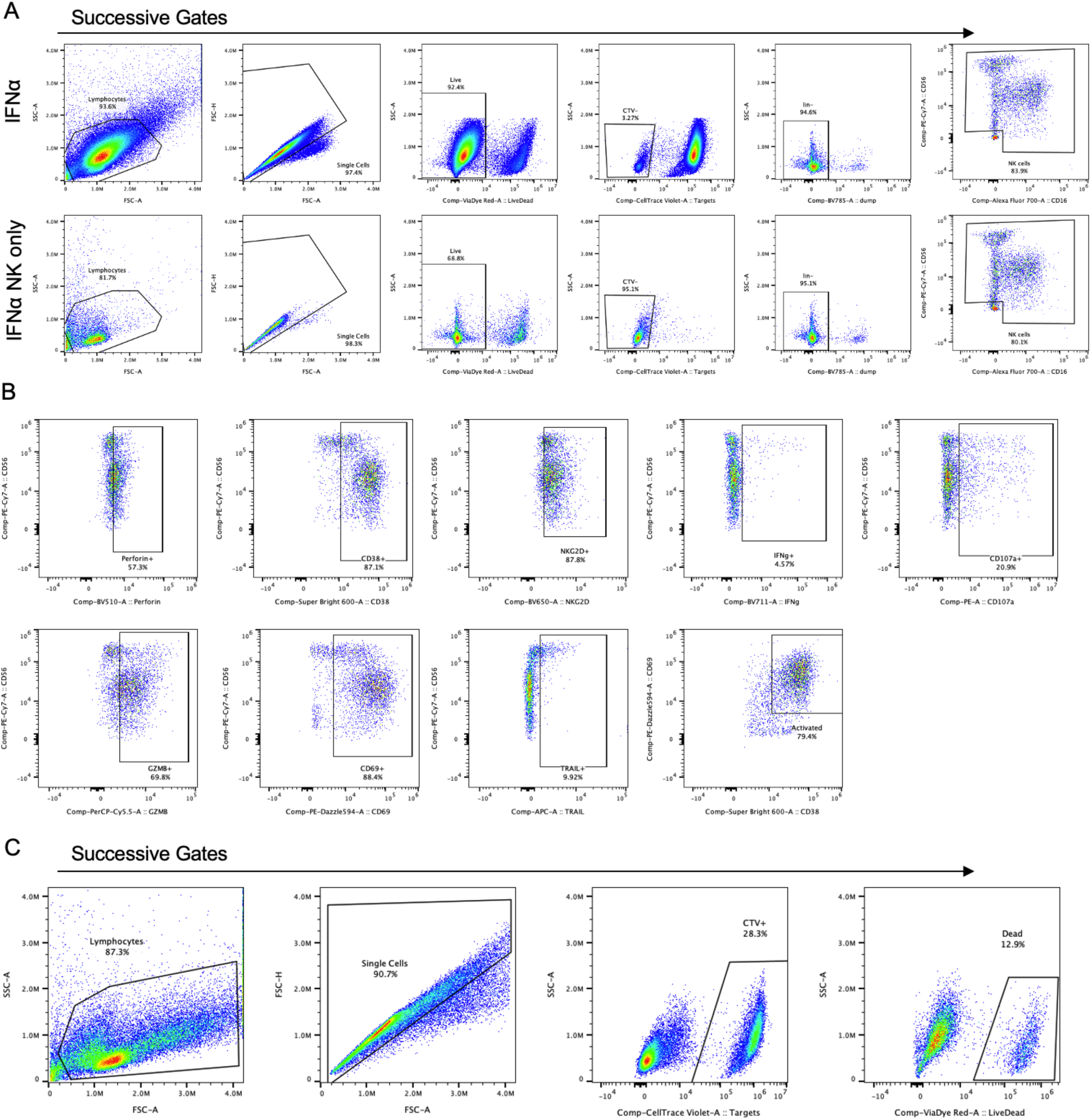
Gating NK Cells and Functional Markers in co-culture with iTfh cells. **A**. Representative Flow plots showing gating scheme to identify NK cells in functional co-culture experiments for one donor in IFNα activated co-culture and mono-culture groups. lin-indicates negative gate for pooled CD3, CD19, and CD14 antibodies and target iTfh-like cells were labeled with cell trace violet (CTV). **B.** Representative flow plot showing expression and gating of functional/phenotypic NK markers in co-culture experiments from one donor in IFNα group. **C.** Representative flow plot showing gating to identify dead target cells in killing assay.

## Supplementary Information

**Supplementary Table 1**

De-identified clinical and demographic metadata for each participant in scRNA-seq and CyTOF data.

**Supplementary Table 2**

NT50 for each patient against each variant from pseudovirus neutralization assays. Each NT50 is the average of 3 technical replicates

**Supplementary Table 3**

Significant differentially expressed genes for each cell type between broad and narrow neutralizers by Seurat’s FindMarkers() using wilcoxon rank sum test, log_2_FC > .25 and adjusted P-value < .05. Significantly Enriched KEGG pathways for each celltype for genes upregulated in broad or narrow neutralizers. Genes Significantly correlated with breadth score for whole PBMCs and NK cells using pseudobulk correlation analysis with DESeq2. Sheet names indicate cell types and breadth groups.

**Supplementary Table 4**

Multinichenetr output of top 50 receptor-ligand interactions show in in Figure 6

**Supplementary Table 5**

MFI and/or percent positivity for markers plotted from in vitro flow cytometry experiments. Processed RT-qPCR data plotted.

**Supplementary Table 6**

Raw RT-pPCR data from experiments plotted.

**Supplementary Table 7**

Antibodies and reagents used for flow cytometry.

**Supplementary Table 8**

Raw luminescence data from pseudovirus neutralization assays against each SARS-CoV-2 variant. Plate layout and key in *Plate Key Columns* and *Plate Key Row* sheets. Sheet names indicate variants used.

## Acknowledgements

We are very grateful to the donors who provided peripheral blood for these experiments and to their families. We thank Drs. Peter S Kim, Jesse Bloom, Payton Weidenbacher, Theodora Bruun, and Duo Xu for the gift of HeLa-ACE2-TMPRSS2 cells, spike plasmids and assistance with pseudovirus neutralization assay protocol. We thank Dr. Dimitra Peppa for her help with co-culture experiments. We thank Kalani Ratnasiri and Dr. Brooks Benard for their input on analysis and visualizations.

This work was supported, in whole or in part, by the Bill & Melinda Gates Foundation OPP1113682. Under the grant conditions of the Foundation, a Creative Commons Attribution 4.0 Generic License has already been assigned to the Author Accepted Manuscript version that may arise from this submission. C.A.B. is an investigator of the Chan Zuckerberg Biohub. This work was also supported by fellowship and training support from National Institutes of Health through Grants T32 AI007290 Molecular and Cellular Immunobiology and SGF: Stanford Graduate Fellowship in Science & Engineering (to I.D.K.).

## Stanford COVID-19 Biobank members

Thanmayi Ranganath, Nancy Q. Zhao, Aaron J. Wilk, Rosemary Vergara, Julia L. McKechnie, Lauren de la Parte, Kathleen Whittle Dantzler, Maureen Ty, Nimish Kathale, Giovanny J. Martínez-Colón, Arjun Rustagi,Geoff Ivison,Ruoxi Pi,Madeline J. Lee,Rachel Brewer,Taylor Hollis,Andrea Baird,Michele Ugur, Michal Tal, Drina Bogusch, Georgie Nahass, Kazim Haider, Kim Quyen Thi Tran, Laura Simpson, Hena Din, Jonasel Roque, Rosen Mann, Iris Chang, Evan Do, Andrea Fernandes, Shu-Chen Lyu, Wenming Zhang, Monali Manohar, James Krempski, Anita Visweswaran, Elizabeth J. Zudock, Kathryn Jee, Komal Kumar, Jennifer A. Newberry, James V. Quinn, Donald Schreiber, Euan A. Ashley, Catherine A. Blish, Andra L. Blomkalns, Kari C. Nadeau, Ruth O’Hara, Angela J. Rogers, Samuel Yang.

